# WNT1 Inducible Signaling Pathway Protein 1 (WISP1) stimulates melanoma cell invasion and metastasis by promoting epithelial – mesenchymal transition

**DOI:** 10.1101/427088

**Authors:** Wentao Deng, Audry Fernandez, Sarah L. McLaughlin, David J. Klinke

## Abstract

Besides intrinsic changes, malignant cells release soluble signals to reshape their microenvironment. Among the signaling factors is WNT1 inducible signaling pathway protein 1 (WISP1), a secreted matricellular protein that is elevated in a variety of cancers including melanoma and is associated with reduced overall survival of patients diagnosed with primary melanoma. In this work, we found that *WISP1* knockout both increased cell proliferation and repressed wound healing, migration and invasion of mouse and human melanoma cells in an ensemble of *in vitro* assays. *In vivo* metastasis assays showed that WISP1 knockout repressed tumor metastasis in both C57BL/6Ncrl and NOD-scid IL2Rgammanull (NSG) mice with B16F10 and YUMM1.7 melanoma cells. Mechanistically, B16F10 cells that invaded in a transwell assay possessed a gene expression signature similar to Epithelial - Mesenchymal Transition (EMT), including coincident repression of E-cadherin and induction of fibronectin and N-cadherin. Upon WISP1 knockout, these EMT signature genes went in opposite directions in both mouse and human cell lines and were rescued by media containing WISP1 or recombinant WISP1 protein. *In vivo,* metastasis repression by WISP1 knockout was reversed by the reintroduction of either WISP1 or SNAI1. A set of EMT gene activation and inhibition experiments using recombinant WISP1 or kinase inhibitors in B16F10 and YUMM1.7 cells suggested that WISP1 activates Akt and MAP kinase signaling pathways to shift melanoma cells from a proliferative to invasive phenotype. Collectively, the results supported a model that WISP1 within the tumor microenvironment stimulates melanoma invasion and metastasis by promoting an EMT-like process.

## INTRODUCTION

Tumor metastasis is a multistep cascade that starts with local invasion into the surrounding tissue and culminates in colonizing distant tissues (1, 2). Classically, melanoma is thought to progress linearly whereby the growth of the primary tumor progressively increases the propensity for metastasis (3). Yet, 4–12% of patients with metastatic melanoma do not have a clinically identifiable primary tumor, and the excised primary melanoma can still recur at different sites in the body as metastatic lesions (4). Observed early dissemination and metastasis of melanoma suggest a more complex, parallel progression model of metastasis in melanoma (4). The basis for this parallel progression model is attributed to reversible phenotype switching of melanoma between proliferative and invasive phenotypes, and the resulting intratumoral heterogeneity, driven by oncogenic signaling and environmental cues (5, 6).

The switch in malignant melanocytes between proliferative and invasive phenotypes resembles the epithelial-mesenchymal transition (EMT), a key process of phenotypic change that is associated with the metastatic progression of epithelial cancers through the control of EMT-inducing transcription factors (EMT-TFs) such as SNAI1/2, ZEB1/2 and TWIST (5, 6). While the specific EMT-TFs that control the phenotypic state depend on cellular context (7–9), this core network is regulated by various signaling pathways that integrate information from environmental cues, including TGF-β, FGF, EGF, HGF, NF-kB, Wnt/β-catenin and Notch pathways (4–6, 10). Of these signaling pathways, genetically engineered mouse models and samples from melanoma patients provide strong evidence on the essential role of Wnt/β-catenin pathway for melanoma development, phenotype switching/EMT, metastasis and drug resistance (4, 11, 12). While it is generally accepted that altered β-catenin signaling is critical for melanoma initiation and proliferation, conflicting roles of β-catenin have been reported for melanoma metastasis (4, 11). Using *Braf^V600E^/Pten^−/−^* and *Braf^V600E^/Pten^−/−^/CAT-STA* mice as melanoma models (13, 14), Damsky et al. found β-catenin activation substantially increased melanoma lung metastasis (14), and Spranger and Gajewski revealed that melanoma-intrinsic active Wnt/β-catenin signaling prevented anti-tumor immunity via T-cell exclusion, thus facilitated tumor progression and metastasis (15). On the other hand, using the *Braf^V600E^/Cdk2a^−/−^/Pten^−/−^* mouse-derived YUMM1.7 melanoma cell line, Kaur et al. discovered that a fibroblast-secreted Wnt antagonist, sFRP2, increased tumor metastasis by repressing β-catenin activity and the expression of MITF, the melanoma differentiation marker microphthalmia-associated transcription factor (16).

Propagation of environmental cues initiated by aberrant signaling within malignant cells, like β-catenin, to reshape the tissue microenvironment is important yet poorly understood (17). Interestingly, β-catenin activation induces a variety of secreted Wnt/β-catenin signaling effectors, including WNT1 inducible signaling pathway protein 1 (WISP1/CCN4) (18–20). As a member of CCN family, WISP1 is a cysteine-rich, secreted matricellular protein that can exert paracrine action by activating PI3K/Akt signaling in fibroblast, cardiomyocyte and neuronal cells, as well as human esophageal squamous cell carcinoma, lung carcinoma and breast cancer cells (21-25). Other signaling activation by WISP1 such as FAK, MEK/ERK, and NF-kB is also reported (26, 27).

In humans, elevated WISP1 expression correlates with poor prognosis in the majority of cancers studied, and WISP1 can promote tumor cell proliferation, survival, and migration/ invasion *in vitro* and tumor growth and metastasis *in vivo* in a variety of human tumors such as breast, pancreatic, prostate, lung, colorectal, and brain cancers (28, 29). For its role during tumor metastasis, recombinant mouse Wisp1 was shown to induce EMT and migration through regulating EMT marker genes (30), while an overexpression of WISP1 in breast cancer cell line (MCF-7) was also reported to both induce cell proliferation and EMT to promote tumor cell migration and invasion (31). However, despite reports in other cancers, the proposed role of WISP1 in melanoma appears to be contradicted and an intracellular signaling basis for these observations remains unclear (18, 32, 33).

Recently, we showed that WISP1 from melanoma cells contributed to tumor immunosuppression (34) and that WISP1 expression correlated with tumor cell invasion in both melanoma and breast cancer (34, 35). Furthermore, disruption of adherens junctions induced the synthesis and release of WISP1 via non-canonical activation of β-catenin (36). Taken together, these findings led us to investigate whether WISP1 is a paracrine effector of Wnt/β-catenin signaling that coordinates EMT/ phenotype switching and metastasis. To test this idea, *in vitro* assays established that WISP1 regulated the growth, migration and invasion of mouse and human melanoma cells. *In vivo* metastasis assays using C57BL/6Ncrl and NOD-scid IL2Rgamma^null^ (NSG) mice suggested a functional role of WISP1 in promoting tumor metastasis, which was supported by retrospective analysis of human transcriptomic, proteomic, and genomic data. Mechanistically, B16F10 cells that invaded Boyden chamber transwells possessed the canonical gene expression profile for EMT including Snail activation and E-cadherin repression. With WISP1 knockout in mouse and human melanoma cells, those EMT markers went to opposite directions and were rescued by conditioned media containing WISP1. *In vivo,* metastasis repression by Wisp1 knockout was reversed by the reintroduction of either Wisp1 or Snail. Collectively, the results supported a model that upregulation of WISP1 within the tumor microenvironment stimulates melanoma invasion and metastasis by promoting an EMT-like process.

## RESULTS

### WISP1 expression is increased in primary melanoma and is associated with reduced overall survival of patients diagnosed with primary melanoma

To ground our study clinically, we first reviewed WISP1 expression in human samples obtained from patients diagnosed with primary melanoma deposited into public databases. Analysis of gene expression profiles reported by Talantov et al. (37) showed *WISP1* mRNA level was almost doubled in primary melanoma samples as compared to benign melanocytic skin nevi (p-value < 0.0002, Fig. 1A). Expression of WISP1 mRNA was not significantly different in benign melanocytic skin relative to normal skin (p-value = 0.795). At the protein level, an independent tissue microarray containing samples from normal epithelial tissue (n= 3) and primary melanoma (n =7) tissue were used to characterize WISP1 expression. Signal deconvolution and image segmentation were used to quantify differences in WISP1 staining in melanocytes and other cells present among architectural features of the skin. In both primary melanoma and normal skin, the protein is located in the cytoplasm (Fig. 1B). In melanoma samples, almost all tumor cells (>75%) exhibited medium to high WISP1 intensity (Fig. 1B, right), while in normal skin there was little or no WISP1 in epidermal keratinocytes, but medium WISP1 staining in melanocytes within both basal layer of epidermis and hair follicles (Fig. 1B, left). Medium WISP1 expression was observed in the fibroblasts in skin dermis (stroma) as well (Fig. 1B, left). The average intensity of WISP1 staining within a tissue sample suggested that an increase in WISP1 also correlates with oncogenic transformation (Fig. 1C, p-value < 0.005). While an increase in average intensity could be explained by a change in cellular composition of the tissue sample, a quantitative analysis of the intensity of WISP1 staining suggests that more of the tissue area stains positive for WISP1, which suggests that more WISP1-producing cells are present, and the staining intensity is greater in primary melanoma than normal skin, which suggests that WISP1-positive cells are producing more WISP1 (Fig. 1D). The results from this quantitative IHC analysis are consistent with the mRNA data presented in Figure 1A such that WISP1 expression was increased in primary melanoma compared to benign skin conditions.

**Figure 1.**
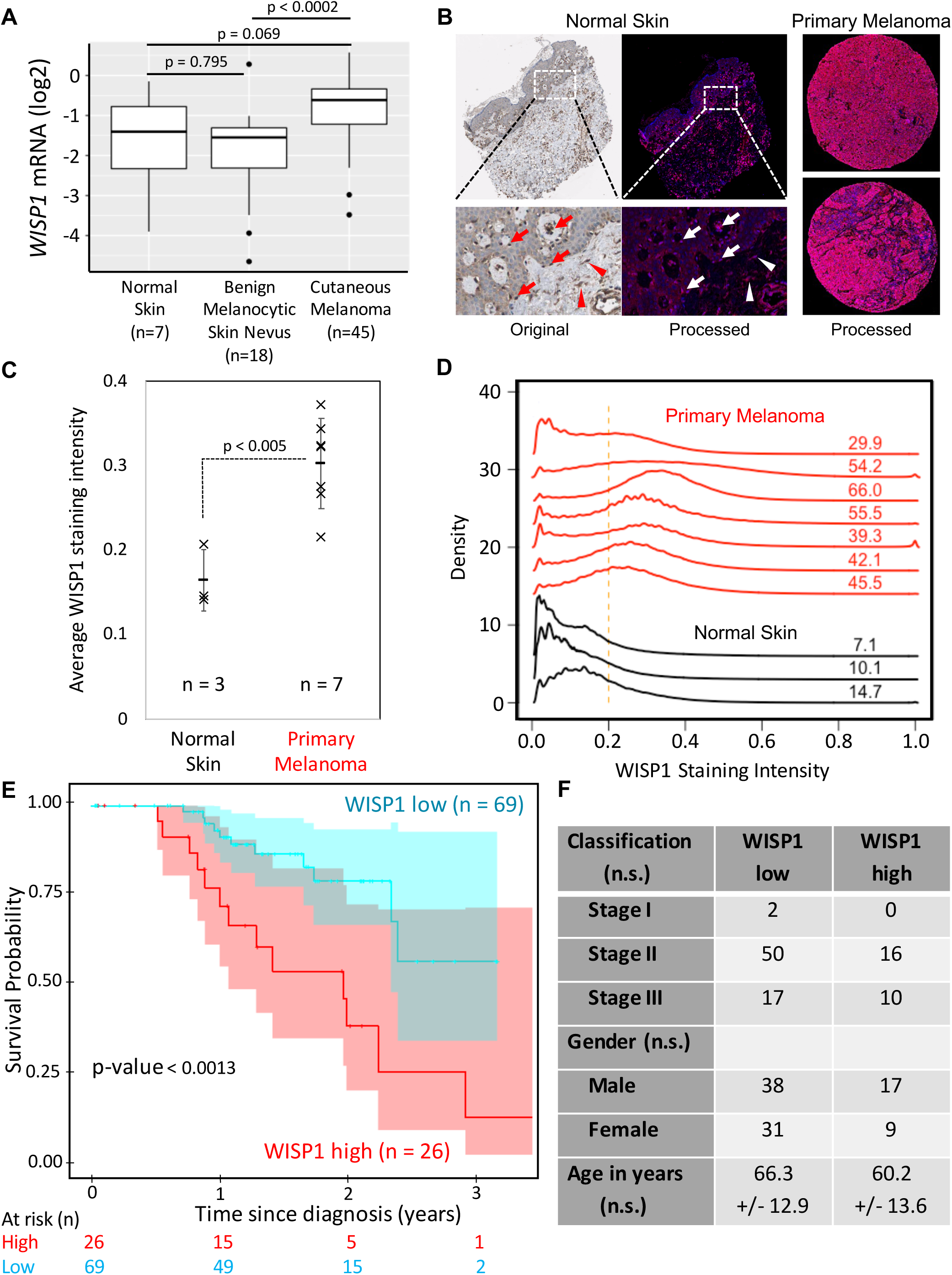
WISP1 expression is increased in melanoma and is associated with reduced overall survival of patients diagnosed with primary melanoma. **A,** Comparison of WISP1 mRNA expression in benign skin conditions (normal skin and benign melanocytic skin nevus) to primary melanoma. Original expression profiles were from (37). P-values calculated using ANOVA with post-hoc Tukey HSD test. **B,** Representative original and deconvoluted color images derived from human normal skin and melanoma tissue microarray probed using a WISP1 antibody (HPA007121) and imaged using 3,3’ diaminobenzidine and stained using hematoxylin for a normal skin (left) and two melanoma (right) tissue samples (original tissue microarray images were obtained from www.proteinatlas.org) (59). Deconvoluted intensity of WISP1 staining is shown in red while cellular structures stained using hematoxylin are shown in blue. Arrows indicate melanocytes in epidermis and arrowheads indicate fibroblasts in dermis (stroma). **C,** The average WISP1 staining within normal skin and primary melanoma tissue samples. **D,** Distributions in non-zero pixel intensity values of WISP1 staining for normal skin (black curves) and primary melanoma (red curves) tissue samples. Numbers indicate the percentage of the distribution that have normalized pixel intensity values greater than 0.2. **E,** Kaplan-Meier estimate of overall survival of patients diagnosed with primary melanoma stratified by WISP1 transcript abundance (data from TCGA). Sample numbers and p-values calculated using the Peto & Peto modification of the Gehan-Wilcoxon test are indicated. **F,** Patient population characteristics of WISP1 high and WISP1 low groups. Statistical differences among categorical data and age were assessed using Fisher’s Exact test and Student’s t-Test, respectively (n.s. indicates p-value > 0.05).

As the IHC and gene expression analyses suggest that malignant transformation of melanocytes is associated with an increase in WISP1 production, we explored genetic mutations, including both coding sequence changes and changes in copy number via structural alterations, that are enriched in melanoma. As WISP1 expression can be induced by non-canonical β-catenin signaling resulting from dynamic turnover of adherens junctions (36), we found that mutations associated with melanoma suggest enhanced malignant cell production of WISP1 (Supplementary Table S1). While mutations in *BRAF, NRAS,* and *CDKN2A,* are highly prevalent in melanoma, mutations also frequently occur in *PTEN, TP53, MITF, KIT, CTNNB1,* and *WISP1*. Notably, the mutation rate (8%) for *WISP1* in melanoma is equal to or higher than those for *CTNNB1, MITF,* and *KIT,* which are considered as promising therapeutic targets (4, 38, 39). Mutations impacting either *WISP1* or *CTNNB1* comprised 13% of the samples. We also noted that mutations in *WISP1* were mainly copy number amplifications (q-value = 2.9E-5) and in *CTNNB1* were mainly single nucleotide variants associated with exon 3. Mutations in exon 3 of *CTNNB1,* which inhibits proteasomal degradation, and copy number amplifications of *WISP1* both favor increased transcriptional response to dynamic turnover of adherens junctions.

To assess the clinical implications of WISP1 overexpression, overall survival of patients diagnosed with primary melanoma stratified by WISP1 expression was estimated using RNA-seq data obtained from 95 patient samples from the TCGA with corresponding survival data. Kaplan-Meier analysis was performed by separating the population into two groups based upon a WISP1 expression cutoff of 1.0 FPKM (WISP1 low n =69, WISP1 high n=26) (Fig. 1E and 1F). While the staging, age, and gender profiles of these two groups are not statistically different, the WISP1 high group patients have a lower 3-year survival rate of only 14% compared to a rate of 58% in the WISP1 low group (p-value<0.0013). The median survival time is about 44 months for WISP1 low group patients, but only 24 months for WISP1 high group patients (Fig. 1E). In addition, a multivariate Cox proportional hazards regression analysis using WISP1 classification (high versus low), tumor stage, and gender as potential covariates of overall survival as the outcome variable indicated that WISP1 classification was the only covariate with a significant association with overall survival (Likelihood ratio test p-value = 0.0061) such that a low value of WISP1 (FPKM < 1) reduces the risk of death by a factor of 0.287 (95% CI: 0.1279 – 0.6442). Collectively, these results suggested that WISP1 expression was elevated in human melanomas and that WISP1 may potentially serve as a biomarker for worse patient prognosis and survival. However as other genes are co-amplified in conjunction with WISP1, we decided to explore the functional impact of WISP1 on melanoma cells to identify a mechanistic underpinning for this difference in patient survival.

### WISP1 knockout in mouse/human melanoma inhibited tumor cell migration and invasion

Using B16 mouse melanoma cell models, we previously reported a role for WISP1 in immunosuppression and its synthesis and secretion following β-catenin release from adherens junctions (34, 36). Following from these studies, ELISA revealed that Wisp1 was secreted into media conditioned in 2D culture by non-metastatic B16F0 cells (605±15pg/ml) and metastatic B16F10 cells (1,300±35pg/ml), as well as immortalized melanocyte Melan-A cells (1018±32pg/ml) (Supplementary Table S2). In comparison, the mouse fibroblast cell line NIH3T3 expressed lower Wisp1 (83±2.2pg/ml), while Wisp1 was almost undetectable (<20pg/ml) in media conditioned by another two mouse tumor cells: Lewis lung carcinoma LLC1 cells and breast cancer E0771 cells (Supplementary Table S2).

To investigate the roles of WISP1 in melanoma progression and metastasis, we knocked out the *Wisp1* gene in metastatic B16F10 cells and evaluated the phenotype of the resulting cell lines using an ensemble of *in vitro* assays that capture aspects of metastasis including cell-matrix and cell-cell interactions, migration and invasion. Using two CRISPR/Cas9 (double nickase) systems to target *Wisp1* gene at two different locations, we cloned two *Wisp1* knockout cells from B16F10 (Supplementary Table S2). In 2D culture, the knockout cells outgrew the parental cells by 96.7±2.5% (F10-KO1) and 73.1±9.5% (F10-KO2) in a two-day period (Fig. 2A). In 3D culture, wild-type (wt) B16F10 and Wisp1 KO cells were used to evaluate the effect of WISP1 on anchorage-independent growth (Fig. 2B). A soft agar assay showed that *Wisp1* knockout in B16F10 cells increased its colony formation by 78.7±6.7% (F10-KO1) and 66.9±7.0% (F10-KO2), respectively. *In vivo, Wisp1* knockout did not affect subcutaneous growth of tumors in NSG mice (Supplementary Fig S2A). In addition, a wound healing assay was used to test the effect of *Wisp1* knockout on tissue repair *in vitro.* For B16F10 cells, the wound healing rate was reduced from 96.3±0.7% (F10) to 61.4±1.4% (F10-KO1) and 65.9±1.0% (F10-KO2) (Fig. 2C). These *in vitro* results suggested that reducing Wisp1 expression increased melanoma proliferation but repressed tumor cell migration, which was consistent with the previous reports that WISP1 repressed melanoma growth both in cell culture and in a mouse model (18, 33).

**Figure 2.**
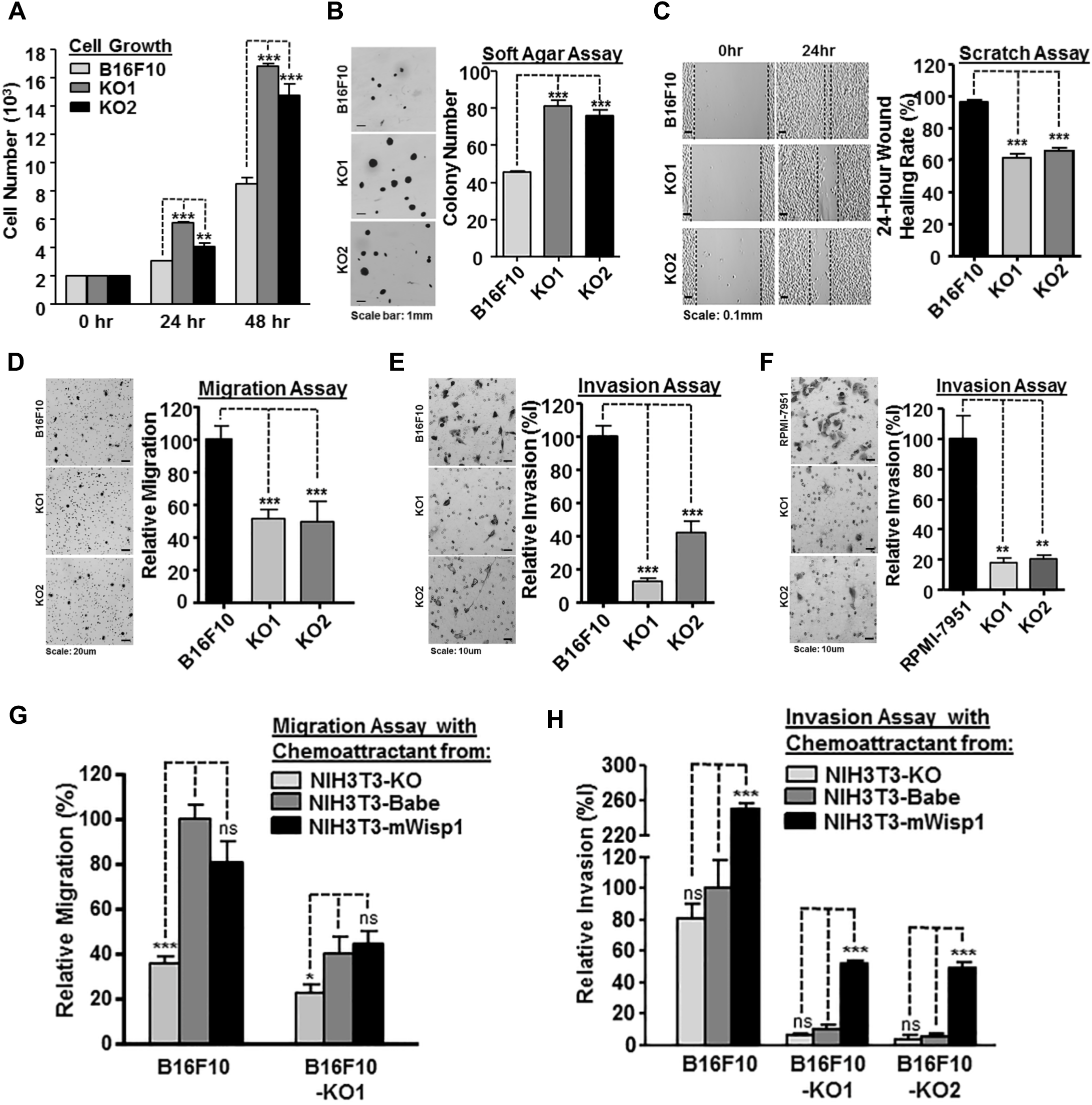
WISP1 knockout in mouse and human melanoma cells inhibited tumor cell migration and invasion. **A,** 48-hour 2D growth of mouse metastatic melanoma cell line B16F10 and two B16F10 Wisp1-knockout cells (-KO1 and -KO2). **B,** Anchorage-independent growth assay of B16F10 and the two knockout cells in soft agar. Colonies were fixed and counted after 14 days. A representative staining image for each sample is shown on left, colony counts is plotted on the right. **C,** Wound healing assay of B16F10 and the two knockout cells. Scratches were created on 6-well plates in biological triplicate and the healing rate was calculated after 24 hours. **D,** Boyden transwell migration assay of B16F10 and the two knockout cells. A representative staining image for each sample is shown on left, relative migration efficiency is graphed on the right. **E,** Boyden transwell invasion assay of B16F10 and the two knockout cells. **F,** Boyden transwell invasion assay of human metastatic melanoma cell line RPMI-7951 and its two Wisp1-knockout cells (-KO1 and -KO2). **G,** Transwell migration assay of B16F10 and its knockout cell (-KO1) using conditioned media with different concentration of Wisp1 as chemoattractant. B16F10 migrated cells with conditioned medium from NIH3T3-Babe were set up as 100% of relative migration efficiency and compared with other cells. **H,** Transwell invasion assay of B16F10 and the two knockout cells using conditioned media with different concentration of Wisp1 as chemoattractant. B16F10 invaded cells with conditioned medium from NIH3T3-Babe were set up as 100% of relative invasion efficiency and compared with other cells. Statistical significance was determined by Student’s t test, where a p-value < 0.05 was considered significant and asterisks was used to indicate calculated range in p-values. *: p-value < 0.05; **: p-value < 0.01; ***: p-value < 0.001; and ns: not significant.

We used Boyden chamber Transwell assays to characterize the effects of WISP1 on both mouse and human melanoma cells. For B16F10, the migration rate of knockout cells was only 51.4±3.2% (F10-KO1) and 49.3±7.4% (F10-KO2) as compared to the parental cells (Fig. 2D), and the invasion rate was reduced even lower to 12.5±1.4% (F10-KO1) and 41.7±7.1% (F10-KO2), relative to the parental cells (Fig. 2E). Among several human melanoma lines we obtained, including RPMI-7951, SH-4, SK-MEL-3 and SK-MEL-24, WISP1 was detected only in medium from RPMI-7951 cells (1,331±34pg/ml) (Supplementary Table S2). After we knocked out *WISP1* in RPMI-7951 using similar CRISPR/Cas9 (double nickase) systems as described above (Supplementary Table S2), we found the invasion rate was repressed significantly to 18.1±2.6% (RPMI-7951-KO1) and 20.6±2.2% (RPMI-7951-KO2), as compared to the parental cells (Fig. 2F).

As illustrated by the tissue microarray IHC results, secretion of WISP1 may also come from tumor stromal cells, such as cancer associated fibroblasts (28, 29, 40, 41). Interestingly, different from most other tumor cells that can use culture medium as a chemoattractant, melanoma cells need conditioned media from mouse fibroblast NIH3T3 cells as a chemoattractant for migration and invasion in the *in vitro* transwell assays (Fig. 2D-2F). As WISP1 was also present in NIH3T3-conditioned media, we next asked if paracrine WISP1 would affect melanoma cell behavior in our assay systems by creating three variants of NIH3T3 cells that had *Wisp1* knocked out using a CRISPR/Cas9 construct, *Wisp1* overexpressed using a retrovirus, and been transduced using a control retrovirus. The knockout cell NIH3T3-KO secreted no detectable WISP1, and the control cell NIH3T3-pBabe provided similar concentration of WISP1 compared to parental cells (90.3±2.8pg/ml) (Supplementary Table S2). The overexpressing cell line NIH3T3-mWisp1 produced about 10 times the concentration of mouse WISP1 protein (920±15.6pg /ml) relative to NIH3T3-pBabe, but its concentration was still lower than what we measured in B16F10 cells.

Using the three different conditioned media as chemoattractants, we evaluated the effect of varying levels of Wisp1 below the transwell insert on the migration and invasion of B16F10, B16F10-KO1 and B16F10-KO2 cells (Fig. 2G and 2H). Generally, existing in chemoattractants, Wisp1 dose-dependently increased the migration and invasion of all three cell lines (Fig. 2F and 2G). The presence of Wisp1, rather than its concentration, appeared to be more important in tumor cell migration (Fig. 2F), while the high concentration of Wisp1, from NIH3T3-mWisp1 cell, seemed to be more decisive in promoting tumor cell invasions no matter whether the melanoma cells expressed Wisp1 or not by themselves (Fig. 2G). Collectively, WISP1 exhibited both autocrine and paracrine effects to stimulate the *in vitro* migration and invasion of melanoma cells. In addition, melanoma invasion, compared with its migration property, responded more drastically to the increased concentration of Wisp1 in its microenvironment.

### Wisp1 knockout repressed mouse melanoma metastasis *in vivo*

Given the effect of WISP1 on melanoma cell migration and invasion in vitro, we next tested the *in vivo* effect of WISP1 on mouse melanoma metastasis using an experimental metastasis assay that directly delivers B16F10 cells into the circulation through mouse tail vein injection. Before injection, all B16F10 and *Wisp1* -knockout cells were transduced with lentivirus expressing a codon-optimized luciferase reporter gene *Luc2* to quantify tumor burden *in vivo.* A real time quantitative PCR (qPCR) method was also developed to quantify the number of metastatic tumor cells (with inserted *Luc2* gene) within defined mouse organs such as lung and liver (42). This method enabled us to detect mouse organ tumor load for as low as one tumor cell within a total of 10^4^ tissue cells (42).

To avoid a confounding influence of host immunity on the response to *Wisp1* knockout (34), we used immunodeficient NSG mice for the first sets of experiments (Fig. 3A-3F). After tail vein injection, wild type B16F10 cells disseminated widely and grew rapidly. Bioluminescence imaging showed the main tumor burden was located in the abdomen (Fig. 3A, left), while the metastatic tumor signals at similar location from B16F10-KO1 cells were much weaker (Fig. 3A, right). Upon dissection, each pair of lungs from B16F10-injected mice were covered with dozens of metastatic tumor colonies, compared to clear lungs from mice receiving B16F10-KO1 cells (Fig. 3B). Real time qPCR revealed that there was an average tumor burden of 85.3±4.2 metastatic tumor cells among 10^4^ lung cells for B16F10-injected mice, compared to an average tumor burden of 7.9±1.1, a reduction of more than 90%, for lungs from B16F10-KO1-injected mice (Fig. 3C). However, lung metastases were a small portion of the overall tumor burden. The majority of B16F10 metastases were observed in the abdomen region, including livers (Fig. 3D), intestines (not shown), and kidney (Fig. 3F). In mice injected with B16F10-KO1 cells, metastatic lesions were either reduced in size and number (livers and intestines, Fig. 3D) or not observed (kidneys, Fig. 3F). Real time qPCR revealed that the average tumor burden in livers from B16F10-injected mice was 1141±136 metastatic tumor cells among 10^4^ liver cells, while the average tumor burden was reduced to 370±26, about a 70% repression, for livers from B16F10-KO1-injected mice (Fig. 3E).

**Figure 3.**
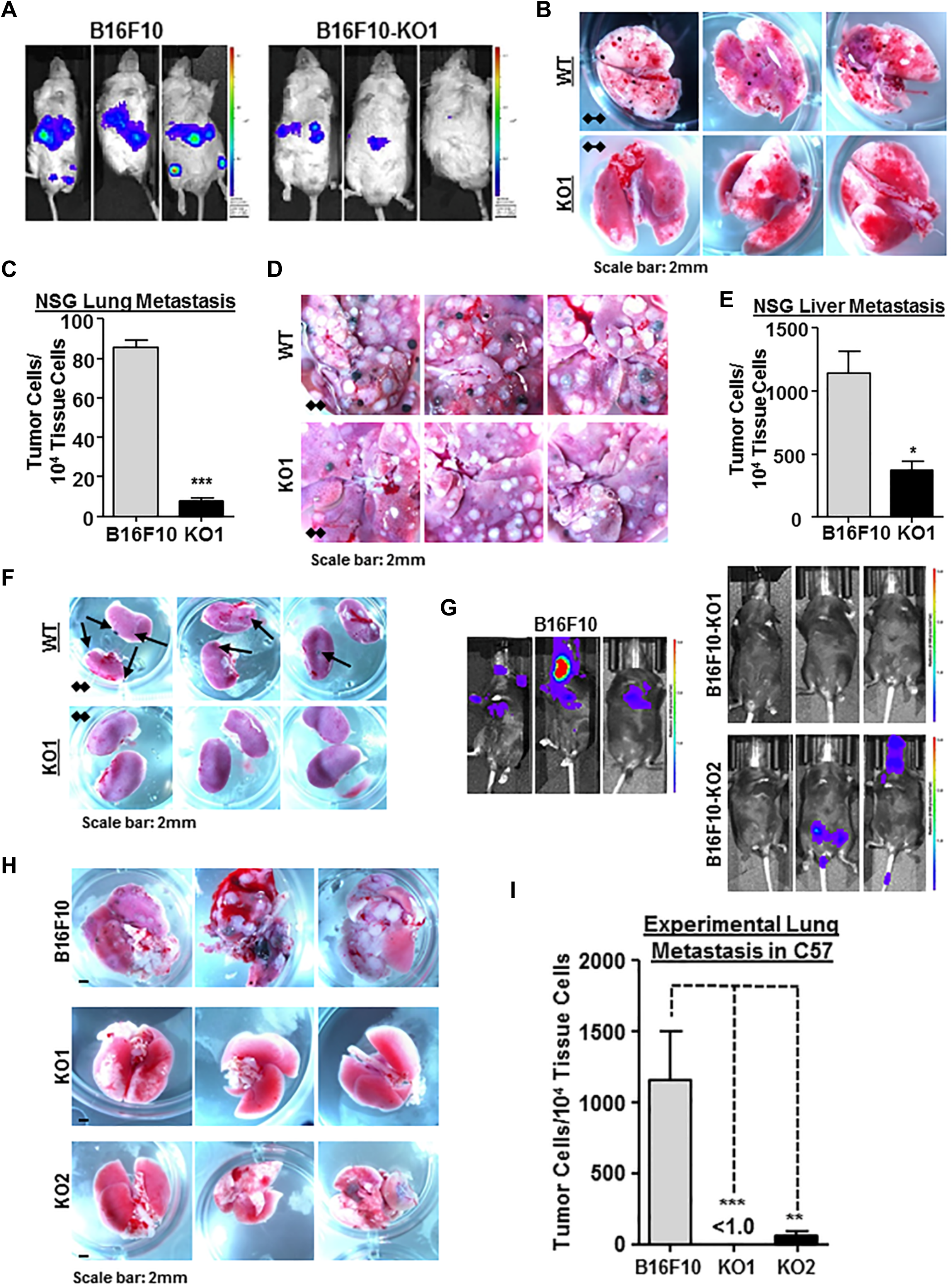
Wisp1 knockout repressed the experimental metastasis of melanoma cell line B16F10 in immunodeficient NSG mice and immunocompetent C57BL/6Ncrl mice. Experimental metastasis assays were performed in NSG mice (**A-F**) and C57BL/6Ncrl mice (**G-I**) using B16F10 and indicated knockout cells with injection through mouse tail veins. Each group contained five duplicates (N=5) and three representative images were shown. These experiments were repeated and similar results were achieved. **A,** Bioluminescence imaging performed one day before NSG mice were euthanized. All animals were compared with the same bioluminescence scale. **B-C,** Tumor lung metastases (black colonies) of NSG mice as captured by photography (**B**) and real time genomic qPCR (**C**). Quantitative tumor lung metastatic burden was assayed and presented as tumor cell number within 10,000 mouse tissue cells. **D-E**, Tumor liver metastases (black and white nodules) of NSG mice as captured by photography (**D**) and real time genomic qPCR (**E**). Quantitative tumor liver metastatic burden was assayed and presented as tumor cell number within 10,000 mouse tissue cells. **F,** Tumor kidney metastases (black colonies) of NSG mice as captured by photography. **G,** Bioluminescence imaging performed one day before C57BL/6Ncrl mice were euthanized. All animals were compared with the same bioluminescence scale. H-I, Tumor lung metastases of C57BL/6Ncrl mice as captured by photography (**H**) and real time genomic qPCR (**I**). *: p-value < 0.05; **: p-value < 0.01; ***: p-value < 0.001.

We next used immunocompetent C57BL/6Ncrl mice for similar experimental metastasis assays (Fig. 3G-3I and Supplementary Fig. S1). After tail vein injection of wild type B16F10 cells, bioluminescence imaging showed that tumor metastases developed in the chest region of all mice (Fig. 3G). In individual mice, signals derived from wt B16F10 cells were also observed in lymph nodes and brains (Fig. 3G, left). In mice injected with one of two *Wisp1* knockout cells (B16F10-KO1 and -KO2), the metastatic tumor signals were consistently absent in the chest region, though individual mice did show signals originating from either the lower abdomen or head (Fig. 3G, middle and right). Dissection confirmed that the majority of tumor metastases associated with wt B16F10 cells were in the lungs (Fig. 3H, upper). In mice injected with *Wisp1* knockout cells, metastatic nodules were either completely absent (B16F10-KO1) or significantly reduced in size and number (B16F10-KO2) (Fig. 3H, middle and lower). Real time qPCR calculated that the average tumor burden in lungs dropped from 1159±349 metastatic tumor cells among 10^4^ lung cells for B16F10-injected mice to less than 1.0 tumor cell (>99.9% reduction) and 57.8±38.7 tumor cells for KO1- and KO2-injected mice, respectively (Fig. 3I). No metastatic nodules were observed on livers from any group of mice (Supplementary Fig. S1A). Although qPCR detected small number of tumor cells in livers, no difference was found between livers from B16F10-injected mice and from B16F10-KO1 mice (Supplementary Fig. S1B).

As experimental metastasis assays suggested that *Wisp1* knockout B16F10 cells had a reduced potential to extravasate and colonize vital organs, we also assessed invasion potential using spontaneous metastasis assays in C57BL/6Ncrl mice with subcutaneous injection of mouse melanoma B16F10 and its *Wisp1-*knockout counterpart (B16F10-KO2). Interestingly, the tumors with *Wisp1* knockout grew slower than wild-type tumors (Supplementary Fig. S2B), which was opposite to the 2D and 3D growth results observed *in vitro* (Fig. 2A and 2B). This difference might be explained by the loss of Wisp1-mediated repression of host anti-tumor immune response, which follows from *in vitro* studies and retrospective analysis of clinical data (34, 35). This hypothesis was supported by the fact that tumors derived from wt B16F10 and WISP1 knockout variants grew at a similar speed in NSG mice (Supplementary Fig. S2A). More focused studies are on-going to clarify this observation. Upon dissection at the humane endpoint of the C57BL/6Ncrl and NSG mice, no metastatic colonies were observed visually in the lungs and liver of mice injected subcutaneously with either wild-type or knockout cells (Supplementary Fig. S2C). However, qPCR surveys of the lungs and livers from C57BL/6Ncrl mice did reveal small micrometastases from B16F10 cells, but nothing from B16F10-KO2 cells (Fig. 4F and Supplementary Fig. S2D).

**Figure 4.**
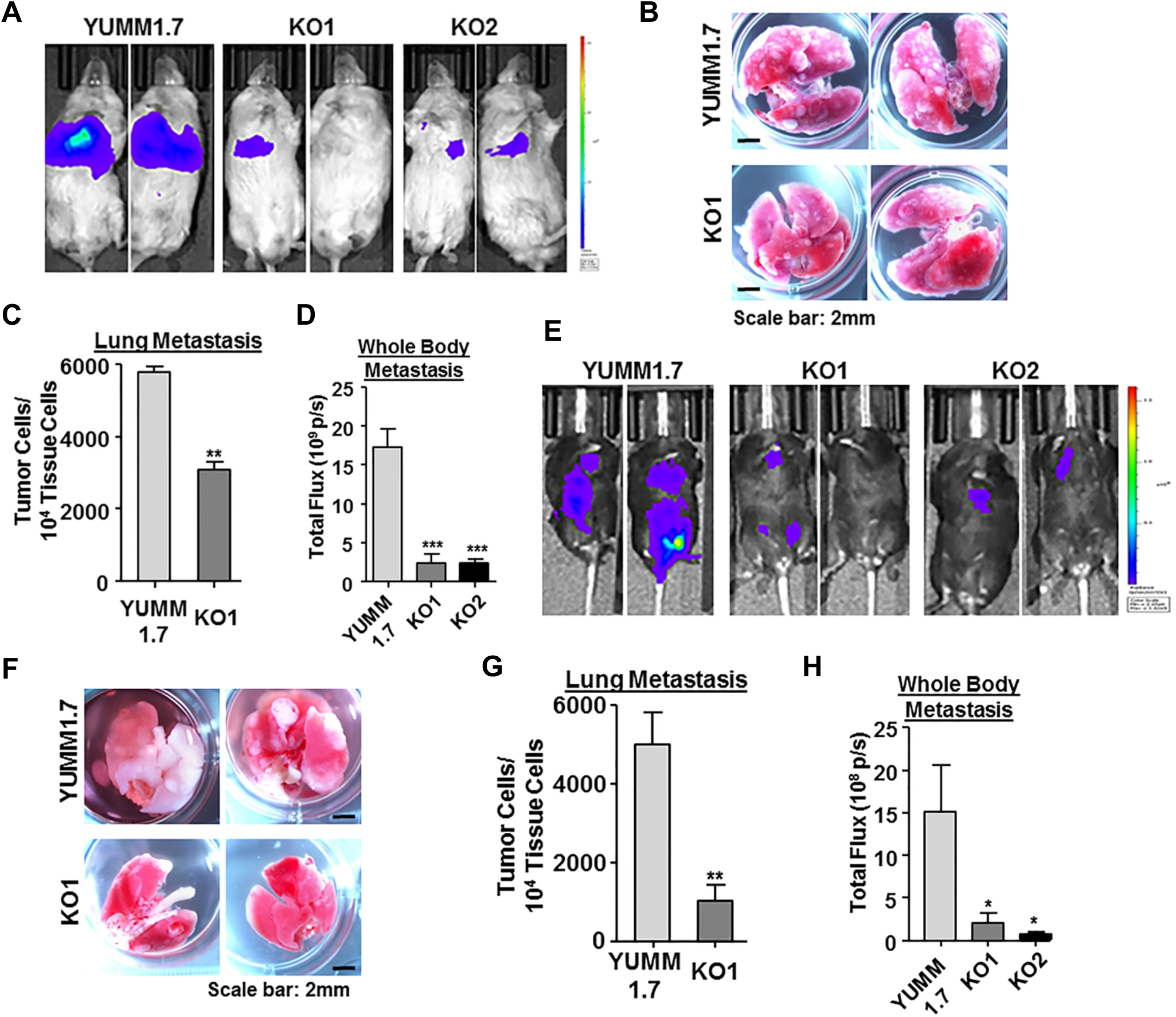
Wisp1 knockout repressed the experimental metastasis of melanoma cell line YUMM1.7 in NSG and C57BL/6Ncrl mice. Experimental metastasis assays were performed in NSG (**A-D**) and C57BL/6Ncrl (**E-H**) mice using YUMM1.7 and indicated knockout cells with injection through mouse tail veins. Each group contained five duplicates (N=5) and two representative images were shown. **A,** Bioluminescence imaging performed one day before NSG mice were euthanized. All animals were compared with the same bioluminescence scale. **B,** Tumor lung metastases (white nodules) of NSG mice as captured by photography. **C,** Real time genomic qPCR quantitatively comparing tumor lung metastatic burdens (tumor cell number within 10,000 mouse tissue cells). **D,** The whole-body metastasis of tumor cells in NSG mice were plotted and compared using bioluminescence intensity detected in panel (A). Total flux is presented as photon/second (p/s). **E,** Bioluminescence imaging performed one day before C57BL/6Ncrl mice were euthanized. All animals were compared with the same bioluminescence scale. **F,** Tumor lung metastases (white nodules) of C57BL/6Ncrl mice as captured by photography. **G,** Real time genomic qPCR quantitatively comparing tumor lung metastatic burdens. **H,** The whole-body metastasis of tumor cells in C57BL/6Ncrl mice were plotted and compared using bioluminescence intensity detected in panel (E). *: p-value < 0.05; **: p-value < 0.01; ***: p-value < 0.001.

B16 variants are relatively unique, chemically induced melanoma models without a defined genetic background and lack BRAF^V600E^ mutations that are prevalent in human melanomas (43, 44). We then tried to reproduce our *in vivo* metastasis assay with more clinically relevant melanoma models. As wt and two WISP1-knockout variants of RPMI-7951 cells failed to survive tail vein injection, we focused next on a series of mouse melanoma cell lines (Yale University Mouse Melanoma, YUMM) recently developed with defined and stable human-relevant driver mutations from genetically engineered C57BL/6 mouse models (44). We tested two of these lines, YUMM1.1 and YUMM1.7 (genotype: BrafV600E/wt Pten-/- Cdkn2-/-), and found YUMM1.7 secreted relatively high amount of Wisp1 (451±25pg/ml), while YUMM1.1 secreted barely detectable Wisp1 in conditioned medium (Supplementary Table S2). We then created two Wisp1-knocked out cells in YUMM1.7 (-KO1 and -KO2) with two sets of CRISPR/Cas9 (double nickase) plasmids and used them in our experimental metastasis assays (Supplementary Table S2 and Fig. 4).

In NSG mice, the main metastatic tumor burden from wild type YUMM1.7 was still located in the abdomen, with much weaker signals at similar location from knockout cells (Fig. 4A). Upon dissection, we found the lungs from YUMM1.7-injected mice were covered with numerous white metastatic tumor nodules, but nodules on the lungs from knockout cell-injected mice were much less and smaller (Fig. 4B). Few visible macrometastatic nodules or colonies were discovered on the liver surfaces from YUMM1.7-injected mice, while nothing visible on the livers from knockout cell-injected mice. This observation is different from B16F10-injected NSG mice, in which liver metastasis from either wild type or knockout B16F10 cells took the majority of overall tumor metastatic burden (Fig. 3D). The quantitative comparison of lung metastasis and whole body metastasis based on real time genomic qPCR and bioluminescence intensity were calculated and plotted in Fig. 4C and 4D.

In C57BL/6Ncrl mice, YUMM1.7 metastasized to a variety of internal organs including lungs, intestines, pancreas, ovary and lymph nodes (Fig. 4E). In mice injected with *Wisp1* knockout cells (-KO1 and -KO2), the metastatic tumor signals were much weaker and detected in much fewer locations in individual mouse (Fig. 4E). The surface of the lungs from knockout cell-injected mice was covered with much less and smaller metastatic white tumor nodules, as compared to YUMM1.7-injected mice (Fig. 4F). Again, the quantitative comparison of lung metastasis and whole body metastasis supported the significance difference observed visually between the wild type cell-injected mice and knockout cell-injected mice (Fig. 4G-H). In general, these *in vivo* results suggest that WISP1 stimulates melanoma metastasis.

### WISP1 stimulated melanoma cell invasion and metastasis through promoting EMT

Following from these *in vitro* and *in vivo* observations, we next focused on identifying a mechanistic basis for how WISP1 promotes a metastatic phenotype. The collective effect of WISP1, one of the Wnt/β-catenin downstream effectors, to inhibit proliferation of melanoma cells while simultaneously promote migration and invasion is reminiscent of EMT-like phenotype switching. In melanoma, the EMT switch starts with the upregulation of EMT-associated transcriptional factors and repression of E-cadherin, as well as the loss of MITF, among other changes in EMT marker genes at the development of melanoma metastasis (5, 6, 39). Therefore, we asked whether an EMT gene signature is influenced by WISP1 and whether EMT-related transcription factors induced by WISP1 regulate tumor cell invasion and metastasis.

Given that the specific genes controlling an EMT switch may depend on cellular context, we first established a gene signature mapped to phenotype by comparing the EMT gene expression profiles between invaded and uninvaded mouse B16F10 melanoma cells in standard Boyden chamber Transwell assays (Fig. 5A). Compared to starting cells and uninvaded cells, the invaded B16F10 cells exhibited a typical EMT gene signature including the activation of EMT transcription factor *Snai1* and *Zeb2,* upregulation of extracellular matrix mesenchymal marker fibronectin (*Fn1*), and downregulation of epithelial marker E-cadherin (*Cdh1*), as well as melanoma differentiation marker *Mitf* (Fig. 5A). The expression of another main EMT transcription factor *Zeb1* was very low and not observed by western (Fig. 5C) in B16F10 cells, which suggested that the observed change in mRNA may not be physiologically relevant. It is unclear whether *Zeb1* plays certain context-specific roles other than promoting EMT, or its function is simply compensated by the redundancy of other EMT-TFs in B16F10 cells. More detailed work is needed to clarify its role associated with B16F10 cell invasion.

**Figure 5.**
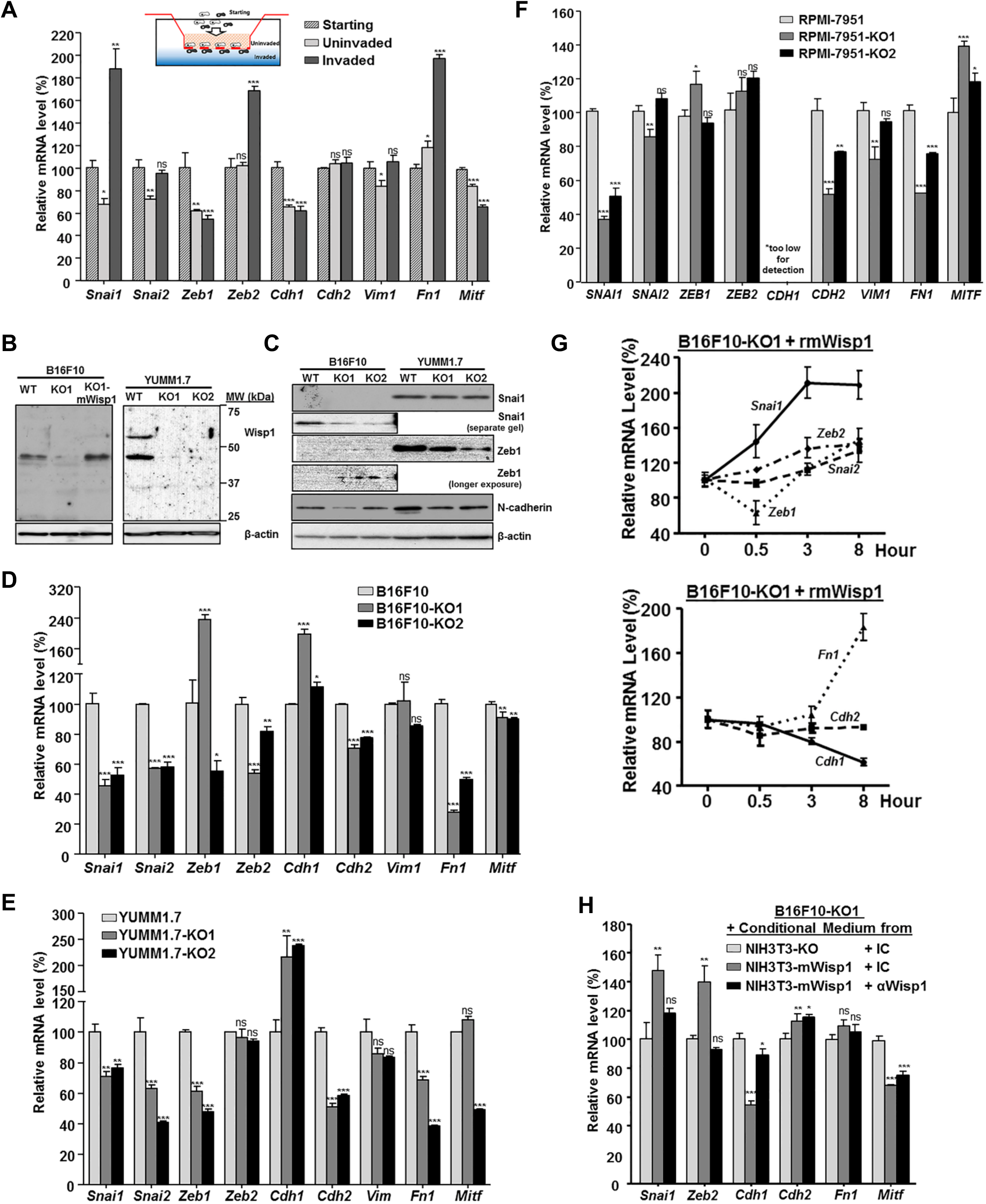
WISP1 induced an EMT gene signature in mouse/human melanoma cells. Unless otherwise specified, all cells were plated on 6-well plates in complete growth medium for 48 hours before harvested for RNA analysis or treated with indicated conditioned medium or recombinant protein. **A,** mRNA expression, revealed by real-time quantitative RT-PCR, of select EMT marker genes and *Mitf* in uninvaded and invaded B16F10 cells from Boyden transwell invasion assay. **B,** Immunoblot analysis of Wisp1 protein to confirm the disruption of Wisp1 gene in B16F10 and YUMM1.7 knockout cells. 20μg of whole cells lysate was load in each lane and P-actin was used as internal loading control. B16F10-KO1-mWisp1 cell, in which mouse Wisp1 expression was resumed with retroviral transduction, was used as a positive control. **C,** Immunoblot analysis of certain EMT marker proteins in B16F10 and YUMM1.7 knockout cells. 20μg of whole cells lysate was load in each lane and all cells were compared on the same gel to reveal the relative intensity of each protein. **D,** Comparison of EMT marker gene expression in mouse melanoma B16F10 and its two Wisp1-knockout cells (-KO1 and -KO2). E Comparison of EMT marker gene expression in mouse melanoma YUMM1.7 and its two Wisp1-knockout cells (-KO1 and - KO2). **F,** Comparison of EMT marker gene expression in human melanoma RPMI-7951 and its two Wisp1-knockout cells (-KO1 and -KO2). **G,** Stimulation of EMT marker gene expression with recombinant mouse Wisp1 protein (rmWisp1). B16F10-KO1 cells were treated with rmWisp1 (final 5μg/ml) and harvested at indicated time point for real-time quantitative RT-PCR analysis. **H,** Stimulation of EMT marker gene expression with Wisp1-overexpressed or Wisp1-immunodepleted conditioned medium. The conditioned media were pre-treated with indicated antibodies for 30 minutes before used on Wisp1-knockout B16F10 cells (-KO1). The cells were collected for real-time qRT-PCR after 3 hour treatment. *: p-value < 0.05; **: p-value < 0.01; ***: p-value < 0.001; ns: not significant.

We then set to compare the expression of genes associated with this EMT signature in *WISP1* -knockout cells to parental cells derived from mouse B16F10, YUMM1.7 and human RPMI-7951 melanoma lines. First, in addition to ELISA (Supplementary Table S2), we used immunoblotting to confirm the knockout of Wisp1 protein in mouse B16F10 and YUMM1.7 cells after the disruption of Wisp1 gene (Fig. 5B). Interestingly, multiple Wisp1 bands were detected in YUMM1.7 cells, suggesting the existence of covalent modification or Wisp1 oligomers. Another set of immunoblotting with available antibodies showed the reduction of EMT transcription factor Snai1 and mesenchymal marker N-cadherin upon *Wisp1* knockout in B16F10 and YUMM1.7 cells (Fig. 5C). While Zeb1 went up in B16F10, it did decrease in YUMM1.7 after *Wisp1* knockout (Fig. 5C).

Using real time qPCR, we found that, in B16F10, those invasion-associated EMT signature genes as determined in Fig. 5A changed in the opposite direction after *Wisp1* knockout (Fig. 5D). The pattern observed upon *WISP1*- knockout is consistent with a Mesenchymal - Epithelial Transition (MET) type switch, which included upregulation of epithelial marker E-cadherin (*Cdh1*), downregulation of EMT-TFs such as *Snai1, Snai2* and *Zeb2,* downregulation of mesenchymal markers such as N-cadherin (*Cdh2*) and fibronectin (*Fn1*) (Fig. 5D). The only exception was the *Mitf* expression from B16F10 knockout cells, which was slightly reduced in both of the B16F10 knockout cells. This may suggest that *Mitf* repression is involved in melanoma cell invasion but is not directly regulated by WISP1 in the context of B16F10 cells. Similar gene expression profiles were discovered in mouse YUMM1.7 and human RPMI-7951 melanoma lines upon *WISP1-* knockout (Fig. 5E-5F). Although we observed subtle differences in the expression of specific genes among the three wild-type and six knockout melanoma cells, WISP1-knockout induced consistently the upregulation of epithelial marker E-cadherin (*Cdh1*), downregulation of EMT transcription factor *Snai1,* downregulation of mesenchymal marker N-cadherin (*Cdh2*) and fibronectin (*Fn1*) (Fig. 5D-5F). The result highly suggested that WISP1 stimulates melanoma invasion and metastasis through promoting EMT of tumor cells, while WISP1 knockout decreases melanoma invasion by causing MET of tumor cells.

A rescue experiment with B16F10-KO1 cells was performed using recombinant mouse Wisp1 protein (rmWisp1) to track the change of these EMT signature genes in real time (Fig. 5H). Within 30 minutes of rmWisp1 treatment, an immediate increase of *Snai1* and decrease of *Zeb1* were observed. Over an eight-hour period, *Snai1* continued to increase and then maintained at a high level, which was followed by the increase of other EMT-TFs, the increase of mesenchymal marker *Fn1,* and the decrease of epithelial marker E-cadherin (*Cdh1*) (Fig. 5H). A similar EMT rescue response in B16F10 knockout cells was observed using conditioned media from mouse fibroblast NIH3T3-mWisp1 that overexpressed mouse *Wisp1* (Fig. 5G, only result from -KO1 shown). Immunodepleting WISP1 in conditioned media prior to treatment of knockout cells abolished such rescue effects (except for *Mitf*), confirming the functional role of WISP1 from the media. Collectively, these results support a notion that WISP1 stimulates tumor invasion and metastasis through promoting an EMT-like process within melanoma cells, and Snai1 plays a major role as a transcription factor in this transition process.

### Snai1 overexpression in *Wisp1*-knockout melanoma cells reversed the repression on tumor invasion *in vitro* and metastasis *in vivo*

The dynamic results described above strongly suggest that SNAI1 is one of main primary effectors downstream of WISP1 signaling to stimulate melanoma cell EMT, hence to promote tumor invasion and metastasis. The idea followed that reintroduction of *SNAI1* into *Wisp1* -knockout melanoma cells would reverse, at least in part, back to the wild-type genotype and phenotype. For such purpose, B16F10 knockout cell was transduced with retroviral vector to overexpress human SNAI1 protein (- KO1-hSnai1) (Fig. 6A). Another two cells from - KO1 were also created either with retroviral vector control (-KO1-pBabe) or with vector overexpressing mouse Wisp1 protein (-KO1-mWisp1) (Fig. 6A).

**Figure 6.**
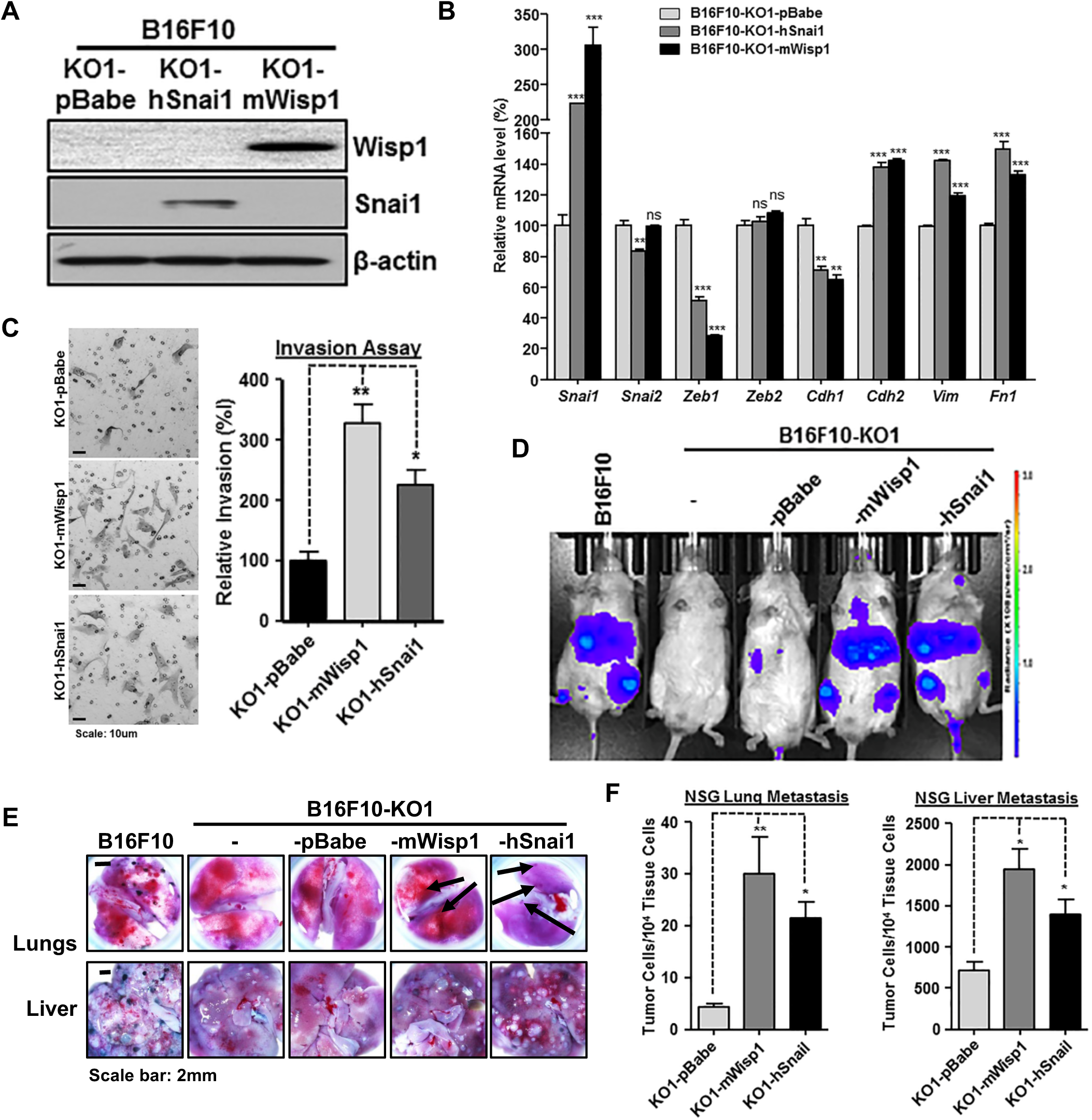
Snai1 overexpression in B16F10 Wisp1-knockout cell rescued the repression on tumor invasion *in vitro* and metastasis *in vivo.* **A,** Immunoblot analysis of Wisp1 and Snai1 using B16F10-KO1 cell that were transduced with retroviral vector control (-pBabe), or retrovirus expressing either mouse Wisp1 (-mWisp1) or human Snai1 (-hSnai1). **B,** Comparison of EMT marker gene expression after overexpression of Snai1 or reintroduction of Wisp1 in B16F10-KO1 cells. Cells were plated on 6-well plates in complete growth medium for 48 hours before harvested for RNA analysis. **C,** Boyden transwell invasion assay after overexpression of Snai1 or reintroduction of Wisp1 in B16F10-KO1 cells. A representative staining image for each sample is shown on left, relative invasion efficiency is graphed on the right. **D,** Experimental metastasis assay in NSG mice using indicated cells. Each group contained 3-4 mice. All mice were imaged one day before the end of the assay and representative bioluminescence images were shown. **E,** Representative lung and liver images from NSG mice in experimental metastasis assay described in panel (**D**). Metastatic tumor colonies on lung surface from mice with (-mWisp1) or (-hSnai1) cells were pointed by arrows. **F,** Real time genomic qPCR for lungs and livers from experimental metastasis assay in panel (**D**). The quantitative tumor metastatic burdens were presented as tumor cell number within 10,000 mouse tissue cells. *: p-value < 0.05; **: p-value < 0.01; ***: p-value < 0.001; ns: not significant.

Real-time qPCR showed that overexpression of either human SNAI1 or mouse Wisp1 in knockout cells recovered the gene expression pattern toward EMT (Fig. 6B). In addition to promoting the expression of endogenous Snai1, they both enhanced the expression of mesenchymal marker N-cadherin (*Cdh2*), vimentin (*Vim*), fibronectin (Fn1), and repressed the epithelial marker E-cadherin (*Cdh1*) (Fig. 6B). The highly similar rescue effects also suggested the two protein factors may work on the same signaling cascades. In transwell assay, -KO1-hSnai1 cell showed an increase in invasion by more than 126.2±25.2% compared to the control cell -KO1-pBabe, and -KO1-mWisp1 cell exhibited an increase of invasion efficiency by over 228.6±29.7% (Fig. 6C).

*In vivo* experimental metastasis assays were performed via intravenous injection using NSG mice. As shown in Fig. 6D, bioluminescence imaging detected very weak metastatic signals in the abdominal region of mice receiving either B16F10-KO1 cells or its retroviral vector control, B16F10-KO1-pBabe cells. Expression of either Wisp1 or Snai1 in knockout cells restored the metastatic phenotype, with a similar intensity as observed with wild type B16F10 cells. Mouse dissection revealed that metastatic tumor colonies on the lung and tumor nodules on the liver, which were absent or significantly reduced after *Wisp1* knockout (Fig. 3B and 3D), were restored upon re-expression of either Wisp1 or Snai1 in knockout cells (Fig. 6E). Real time qPCR confirmed the significant increase in metastasis after Wisp1 or Snai1 were re-expressed in knockout cells (Fig. 6F). In these experiments, the average lung tumor burden was 4.3±0.6 metastatic tumor cells among 10^4^ lung cells for mice receiving -KO1-pBabe cells, the number was increased to 30.0±7.0 and 21.3±3.2 for mice with -KO1-mWisp1 and -KO1-hSnai1 cells, respectively (Fig. 6F, left). Similarly, the average liver tumor burden was 715±110 metastatic tumor cells for mice with -KO1-pBabe cell, and increased to 1944±249 and 1391±186 for mice with -KO1-mWisp1 and -KO1-hSnai1 cells, respectively (Fig. 6F, right). Collectively, these *in vitro* and *in vivo* results supported our proposed role of Snai1 as a downstream effector of Wisp1 signaling and illustrate the role of this signaling pathway in promoting melanoma cell metastasis.

### Wisp1 activated Akt/MAP kinase signaling to promote EMT in mouse melanoma cells

WISP1 activates Akt/PKB signaling pathway to promote cell proliferation and survival in fibroblast, cardiomyocte, neuron and multiple cancer cells (21-25). It also stimulates the MEK/ERK pathway to enhance tumor migration in human chondrosarcoma and osteosarcoma cells (26, 27). Since Akt/PKB signaling and MEK/ERK signaling are the intracellular signaling cascades best known to induce EMT and tumor metastasis (4-6), we hypothesized that these two signaling pathways are essential for EMT in melanoma cells, and that WISP1 activates these signaling pathways to promote EMT.

To test this, we blocked either Akt, MEK, or both pathways in B16F10 cells using kinase inhibitors, and compared the change on EMT marker gene expression after 3 hours (Fig. 7A). As expected, we observed a shift in gene expression toward MET whenever Akt or MEK signaling was inhibited, with at least an additive effect when both were blocked. Changes included the reduction of EMT-TFs such as *Snai1, Snai2, Zeb2,* and the increase of epithelial marker E-cadherin (*Cdh1*) (Fig. 7A). Among the three mesenchymal markers tested, only fibronectin (*Fn1*) showed significant decreases, probably because the 3-hour treatment was not long enough to exhibit any difference on other genes (Fig. 7A). A similar MET pattern was observed in YUMM1.7 cells upon treating with kinase inhibitors (Fig. 7B). We also saw the reduction of *Zeb1,* which was consistent with the EMT gene expression pattern we observed for YUMM1.7 in Fig. 5E.

**Figure 7.**
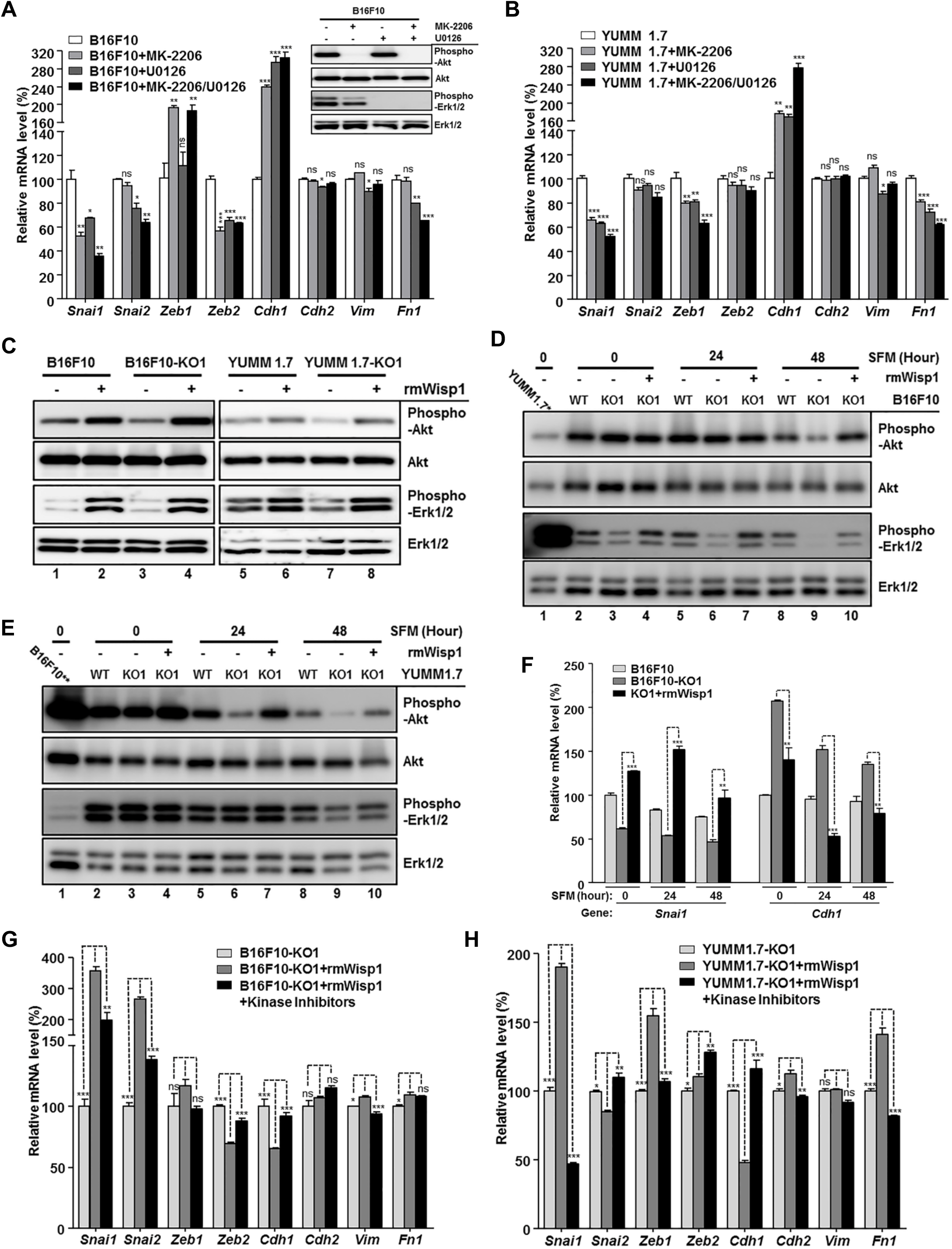
Wisp1 activated Akt/MAPK signaling and promoted EMT marker gene expression in mouse melanoma cells. Unless otherwise specified, cell treatment for kinase immunoblot analysis maintained for 30 minutes before cells were lysed for protein extraction while cells were treated for 3 hours prior to RNA extraction for comparing EMT marker gene expression. **A,** EMT marker gene expression after inhibiting Akt and/or MAPK signaling in B16F10 cells. Cells were treated with specific phospho-Akt inhibitor MK-2206 and/or phospho-MAPK inhibitor U0126. Immunoblot for phospho-Akt and phospho-Erk1/2 inhibition was shown on the right upper corner. Pan-Akt and total Erk1/2 were also probed as loading control. **B,** EMT marker gene expression after inhibiting Akt and/or MAPK signaling in YUMM1.7 cells, with DMSO as control. **C,** Immunoblot for phospho-Akt and phospho-Erk1/2 in indicated mouse melanoma cells with treatment of recombinant mouse Wisp1 protein (rmWisp1, final 5μg/ml). Cells grown on 6-well plates in complete DMEM for 48 hours and serum-free DMEM (SFM) for another 48 hours before rmWISP1 was added. Pan-Akt and total Erk1/2 were probed as loading control. **D,** Immunoblot analysis of Akt/MAPK activation in B16F10 knockout cell (-KO1) by rmWisp1 under different basal phospho-kinase levels. All cells were grown on 6-well plates in complete DMEM for 48 hours (0 hour point for SFM) and switched to SFM for 24 hour or 48 hours. Indicated cells were treated with rmWisp1 for 30 minutes following 0, 24, or 48 hours in SFM before analyzed for kinase activation. The first lane loaded with YUMM 1.7 at 0 hour to compare the relative kinase level between B16F10 and YUMM1.7 cells. **E,** Immunoblot for Akt/MAPK activation in YUMM1.7 knockout cell (-KO1) by rmWisp1 under different basal phospho-kinase levels. All cells were treated similarly as described in panel (D). The first lane loaded with B16F10 at 0 hour to compare the relative kinase level between B16F10 and YUMM1.7 cells. F Snai1 activation and E-cadherin repression in B16F10 knockout cell (-KO1) by rmWisp1 under different basal phospho-kinase levels. All cells were treated similarly as described in panel (D) except that rmWisp1 treatment at each point maintained for 3 hours. G-H, EMT marker gene expression after Akt/MAPK activation in B16F10-KO1 (G) or YUMM1.7-KO1 (H) by rmWisp1 was blocked by Akt/MAPK inhibitors. rmWisp1 with DMSO or inhibitors was added after indicated cells were grown on 6-well plates in complete DMEM for 48 hours and in SFM for 24 hours. *: p-value < 0.05; **: p-value < 0.01; ***: p-value < 0.001; ns: not significant.

To assay gene expression, melanoma cells including B16F10 and YUMM1.7 were normally plated in complete DMEM (10% FBS) for 48 hours before harvested for RNA extraction. However, to detect maximal Akt and MEK signaling activation with minimal background noise from FBS, we grew these melanoma cells (wild-type and knockouts) in serum-free medium (SFM, 0.1% FBS) for another 48 hours before we treated cells with rmWisp1 for 30 minutes and lysed cells for immunoblotting analysis (Fig. 7C). Both B16F10 and YUMM1.7, with autocrine Wisp1 secretion, maintained a higher basic level of phospho-Akt and phospho-MEK than their knockouts (Fig. 7C, compare lane 1 to 3, lane 5 to 7) and paracrine rmWisp1 treatment similarly elevated phospho-Akt and phospho-MEK of both wild type and knockout cells (Fig. 7C, compare lane 2 to 4, lane 6 to 8). Collectively, Wisp1 rapidly and efficiently activated Akt and MEK signaling pathways that were critical signal transducers for inducing an EMT-like gene expression signature in both B16F10 and YUMM1.7 cells.

While these results are consistent with our hypothesis, we noticed some subtle difference in how these two melanoma models responded to different growth conditions. We designed experiments to explore how pre-conditioning in serum-free medium and the presence of the BrafV600E mutation in YUMM1.7 cells impacted the signaling response to Wisp1. In two sets of time-course experiments, we grew the indicated cells in complete DMEM for 48 hours (SFM, 0 hour time point) and switched to SFM for 24 hour or 48 hours. At SFM time point (PI) 0, 24, or 48 hours, cells were treated with rmWisp1 for 30 minutes before assaying kinase activation. Immunoblotting revealed that B16F10 and its knockout cells exhibited relative high phospho-Akt but low phospho-MEK background level (Fig. 7D, compare lane 1 with 2-4), while YUMM1.7 and its knockout cells exhibited relative low phospho-Akt but high phospho-MEK background level (Fig. 7E, compare lane 1 with 2-4). Hence for B16F10-KO1 cell, rmWisp1 readily stimulated MEK signaling at time point 0, but showed stimulation of Akt signaling at a later time (time point 48) (Fig. 7D). Certainly for YUMM1.7-KO1 cell, rmWisp1 mainly stimulated Akt signaling because of their BrafV600E mutation and activated Braf/MEK/ERK pathway (Fig. 7E). Using similar experimental conditions, a time-course gene expression analysis was also performed for *Snai1* and *Cdh1* (Fig. 7F). The result confirmed the effect of rmWisp1 on EMT switch at all three time points and also suggested the best time for maximal stimulation, which was 24 hours in SFM (Fig. 7F). Under this optimal condition, we stimulated B16F10-KO1 and YUMM1.7-KO1 cells with rmWisp1 in the presence or absence of both Akt and MEK inhibitors to test whether activation of Akt and MEK signaling was essential in melanoma cells for a WISP1-mediated EMT switch (Fig. 7G-7H). Inhibiting these two signaling pathways dramatically repressed the elevation of EMT transcription factors including *Snai1* in B16F10-KO1 and *Snai1/Zeb1* in YUMM1.7-KO1 and reversed the repression on *Cdh1* expression from those EMT-TFs (Fig. 7G-7H). Similar to Fig. 7A and 7B, we did not see much change (except for *Fn1* in YUMM1.7-KO1) with the three mesenchymal markers (*Cdh2, Vim, Fn1*), due to short period of treatment time (Fig. 7G-7H). In general, our results strongly supported the notion that Wisp1 promotes melanoma EMT by stimulating Akt and MEK/ERK signaling pathways.

## DISCUSSION

For melanoma invasion and metastasis, revealing factors present within the tumor microenvironment that regulate these processes has important implications on the diagnosis, prognosis and treatment of melanoma. In this work, analysis of molecular and survival data derived from patients diagnosed with primary melanoma showed the expression of Wnt-inducible Signaling Protein 1 (WISP1), a downstream effector of the Wnt/β-catenin pathway, is increased in melanoma and is associated with reduced overall survival of patients diagnosed with primary melanoma. To establish a functional implication of WISP1, we found that WISP1 enhanced tumor invasion and metastasis by promoting melanoma EMT using metastatic mouse and human melanoma cell lines. Results from experimental metastasis assays with both NSG and C57BL/6Ncrl mice injected with either B16F10 or YUMM1.7 melanoma cells supported the functional role of Wisp1 in vivo. Collectively, these observations for WISP1 revealed a connection back to aberrant Wnt/β-catenin signaling and provide insight into the context-dependent role of Wnt/β-catenin pathway in melanoma metastasis.

Clarifying the *in vivo* relations between aberrant Wnt/β-catenin signaling and functional implications of WISP1 signaling may help explain the prevalence of early metastatic dissemination in melanoma. As a secreted signaling molecule, WISP1 connects intrinsic cell signaling pathways with biological cues released into the tissue microenvironment to restore homeostasis following tissue damage (45-47) and to sustain a mesenchymal stem cell niche (48). While these studies focus on bone and cartilage homeostasis, the expression of WISP1 by normal melanocytes and dermal fibroblasts (Fig. 1A-1D) and the induction of WISP1 expression upon disruption of adherens junctions (36) suggest that WISP1 plays a similar role in the skin. Given similarities between stromal-epithelial cross-talk in wounds and tumors (49), it follows then that melanocytes are poised for metastasis, which is realized by acquiring mutations that amplify the production of this environmental cue. This model is supported by the genetic evidence that WISP1 gene amplification was enriched in melanomas and that β-catenin gene amplification and stabilizing exon 3 mutations were also enriched (Supplementary Table S1). Interestingly, the majority of these changes associated with aberrant β-catenin signaling were independent of WISP1 amplification in patient samples (Supplementary Table S1).

Mechanistically, melanoma invasion and metastasis may be connected with an EMT-like process through the upregulation of EMT-related transcription factors and repression of E-cadherin (4–6). Our results showed that WISP1 upregulated a series of EMT transcription factors and mesenchymal markers and also repressed the epithelial marker E-cadherin as well as the melanocyte differentiation marker MITF (Fig. 5-Fig. 7). The regulation of EMT marker genes by WISP1 was achieved, at least partially, through the activation of Akt and MAP kinase signaling pathways in B16F10 and YUMM1.7 melanoma cells (Fig. 7), whereby kinase inhibitor experiments supported the upstream role of these signaling pathways. Identifying pathways that become activated in response to WISP1 is helpful as, while integrin α5β1 and toll-like receptor TLR4 have been suggested (26, 50-52), the membrane-proximal signaling events associated with WISP1 stimulation remain unclear. Our observed activation of PI3K/Akt and MEK/ERK suggests integrin involvement over TLR4 and is consistent with other reports in different cellular contexts (21–27, 50–52) although functional consequences of WISP1 stimulation are not entirely clear from these studies. Here, we observed that WISP1 switched malignant melanocytes from a proliferative to an invasive phenotype. In terms of proliferation, the results presented are consistent with previous reports showing that both recombinant WISP1 and WISP1 overexpression inhibited melanoma growth *in vitro* and *in vivo* (18, 33). In terms of invasion, we report new findings and explore the gene regulatory landscape to probe for a mechanistic basis. Transcriptionally, the observed changes in gene expression were largely conserved between mouse and human cell lines and consistent with conceptual models of EMT, especially with the coincidental reduction in the epithelial marker gene E-cadherin and induction of the mesenchymal marker genes fibronectin and N-cadherin (4–6). One intriguing observation was the different levels of basal expression of *ZEB1* and the change in expression during EMT induction in three melanoma cell lines we tested. When we knocked out *WISP1* in these melanoma cell lines, we found, with decreased *SNAI1* expression in all cells, *ZEB1* was reduced only in YUMM1.7 from a high basal level of expression, barely changed in RPMI-7951, and even increased in B16F10 cells from a low basal level of expression. Such counterintuitive response of certain EMT-related transcription factors have been previously reported during the switch of primary melanoma to a mesenchymal-like invasive phenotype for another EMT-inducing transcription factor, ZEB2 (7–9). While the observed dynamics of EMT transcription factor response to WISP1 suggest that *SNAI1* induction is a primary response and that *ZEB1* appears to be a secondary response, a more systematic analysis of the dynamics of the underlying network of EMT transcription factors under these different conditions may help reveal how environmental and contextual differences collectively influence EMT.

Targeting the Wnt/β-catenin pathway has been proposed to treat human cancers including melanoma, yet a few conceptual and safety concerns challenge developing this therapeutic approach (53-55). Some of these challenges related to β-catenin stem from it being an intracellular target that plays various roles depending on context, such as in cancer progression, organismal development, and adult tissue homeostasis (53). As a secreted downstream effector of aberrant Wnt/β-catenin in melanoma, targeting WISP1 has several advantages. First, targeting WISP1 may provide more specificity in reshaping the melanoma microenvironment to favor anti-tumor immunity and to inhibit metastasis than pleiotropic effects of inhibiting β-catenin. Second, a secreted target opens more options for developing therapeutic reagents, including humanized monoclonal antibodies against WISP1 to block its activities, siRNA and other oligonucleotides to repress WISP1 expression, or small molecules and peptides to inhibit WISP1 signaling. While there is some evidence suggesting receptors for WISP1 (50, 56), clarifying membrane proximal signaling events that lead to activation of EMT-related transcription factors, including SNAI1, may provide additional extracellular targets. Future pre-clinical studies with animal models exhibiting the full spectrum of melanoma progression will be of great interest for translating WISP1 as a target into the clinic to limit metastatic dissemination for patients diagnosed with melanoma.

## EXPERIMENTAL PROCEDURES

### Cell Culture, WISP1 ELISA and Conditioned Medium Preparation

Mouse melanoma line B16F0 (Purchased in 2008, RRID: CVCL_0604), B16F10 (Purchased in 2008, RRID: CVCL_0159), mouse fibroblast line NIH3T3 (Purchased in 2007, RRID: CVCL_0594), mouse Lewis lung carcinoma line LLC1 (Purchased 05/2017, RRID: CVCL_4358), HEK293T (Purchased in 2005, RRID: CVCL_0063), human metastatic melanoma cell lines RPMI-7951 (Purchased 07/2015, RRID: CVCL_1666), SK-MEL-3 (Purchased 07/2015, RRID: CVCL_0550) and SH-4 (Purchased 07/2015, RRID: CVCL_1692) were purchased from American Type Culture Collection (ATCC) on the indicated dates. Mouse breast cancer line E0771 was kindly provided by Dr. Linda Vona-Davis (Received 08/2015, RRID: CVCL_GR23 - West Virginia University). Mouse melanoma lines YUMM1.1 (Received 09/2017, RRID: CVCL_JK10) and YUMM1.7 (Received 09/2017, RRID: CVCL_JK16) were gifts from Drs. William E. Damsky and Marcus W. Bosenberg (Yale University) (44). B16F0, B16F10, NIH3T3, 293T, YUMM1.1 and YUMM1.7 cells were cultured in high-glucose DMEM supplemented with L-Glutamine, Penicillin-Streptomycin and 10% fetal bovine serum (FBS). They may also be cultured in the same medium with only 0.1% FBS (serum-free medium, SFM) as indicated in the text. Other cells were grown as recommended by ATCC. All cells lines were revived from frozen stock, used within 10-15 passages that did not exceed a period of 6 months, and routinely tested for mycoplasma contamination by PCR.

To measure WISP1 secretion from each line, cells were grown for 48 hour to reach about 90% confluence and the media was filtered for ELISA (enzyme-linked immunosorbent assay) analysis using Human WISP-1/CCN4 DuoSet ELISA Development Kit (R&D Systems, Minneapolis, MN). To prepare conditioned media with different concentration of Wisp1, wt and derivative NIH3T3 cells (NIH3T3-KO, - pBabe, and -mWisp1) were counted and seated on 100mm plates with the same density. Cells were grown in DMEM with 0.1% FBS or 10% FBS for 48 hours to reach about 70% confluence. The media was then filtered, aliquoted and frozen at -80°C for future use. Generally, conditional media with 0.1% FBS were used for transwell migration and invasion assays, while conditional media with 10% FBS were used for gene expression stimulation (Fig. 5G).

### Retroviral/Lentiviral Plasmids and Virus Transduction

Mouse Wisp1 DNA sequence encoding the total 367 amino acids was amplified by PCR from B16F0 cDNA using primers with BamH I site on each side. A retroviral expression vector for mouse Wisp1 (pBabe-mWisp1) was created by inserting the above coding sequence into the BamH I site of pBabe-puro retroviral vector, and was verified by sequencing. Another retroviral vector for human Snail (pBabe puro Snail, or pBabe-hSnai1) was from Addgene (Plasmid # 23347, Gift of Bob Weinberg). Lentiviral vector pLU-Luc2, expressing a codon-optimized luciferase reporter gene *Luc2*, was kindly provided by Dr. Alexey V. Ivanov (West Virginia University) and was described previously (42).

Retroviruses were packaged and transduced into indicated cells. The stable cells were achieved with puromycin selection. Lentiviruses were produced by transfecting pLU-Luc2 and two packaging plasmids, psPAX2 (Addgene plasmid #12260) and pCMV-VSG-G (Addgene plasmid #8454), into HEK293T cells. Virus soup was aliquoted and used to transduce indicated cells at constant conditions.

### Creation of *WISP1*-Knockout Cells Using CRISPR/Cas9 System

To achieve high specificity and reduce variability in genetic backgrounds, CRISPR/Double Nickase systems were selected to knock out the *WISP1* gene. Two pairs of mouse Wisp1 Double Nickase Plasmids (sc-423705-NIC and sc-423705-NIC-2), targeting mouse Wisp1 gene at different locations, were purchased from Santa Cruz Biotechnology (Dallas, Texas) and used in B16F10, YUMM1.7 and NIH3T3 cells. Another two sets of WISP1 Double Nickase Plasmids against human WISP1 gene (sc-402559-NIC and sc-402559-NIC-2) were also from Santa Cruz Biotechnology and used in RPMI-7951 cells.

Following manufacturer’s instructions, cells were transfected with individual set of plasmids, which also express a puromycin-resistance gene. Cells were selected by puromycin for five days to achieve 100% transfection efficiency. The survived cells were counted and plated into 96-well plate with a density of 0.5 cell/well. After one week, the grown-up single clones were isolated and expanded on 6-well plates. The cell culture media from those wells were used for WISP1 ELISA to characterize knockout clones. The identified *WISP1* -knockout cells were further expanded and used for the next steps.

### 2D Cell Growth Assay, Soft Agar Assay and Wound Healing Assay

Two-dimensional cell growth was tested on 96-well plates in biological triplicate using ATPlite Luminescence Assay System (Perkin Elmer Inc., Bridgeville, PA) following manufacturer’s instructions. Anchorage-independent cell growth (Soft Agar Assay) was performed on 6-well plates in biological triplicates as described (57).

For wound healing assays, all cells were prepared on 6-well plates in biological triplicates and allowed to reach 95% confluence. A wound in each well was created by scratching straight though the middle of the well with a 200μl pipette tip. Plates were washed to remove dislodged cells and debris, refed with fresh media, and incubated at 37°C for 24 hours. The center of each scratch was photographed at 0 and 24 hour time point, the relative wound width was measured with ImageJ v1.42 (National Institutes of Health), and the healing rate was calculated.

### Transwell Migration, Invasion Assays and Collection of Invaded Cells

BioCoat Control Inserts for migration assay and BioCoat Matrigel Invasion Chambers for invasion assay were from Corning Inc. (Corning, NY). The assays were performed on 24-well plates in biological triplicates following manufacturer’s instructions. Briefly, cells were serum-starved for 24 hours before trypsinized and resuspended in DMEM with 0.5% BSA. Each well was filled with 0.75ml of serum-free conditioned media from either NIH3T3 or other indicated cell as chemoattractant. The chamber inserts were then placed onto wells and 5.0X10^4^ cells in 0.5ml suspension were loaded into the inserts. The plates were incubated at 37°C for 24 hours, the cells on the upper surface of the insert PET membrane were carefully removed with a cotton swab and the cells that migrated or invaded through the membrane were stained with Hema 3 Staining System (Thermo Fisher Scientific Inc., Waltham, MA). The membrane was peeled off with a razor blade and mounted on a glass slide. The cells were then quantified by microscope.

To collect uninvaded and invaded B16F10 cells in the transwell assay for RNA isolation and gene expression analysis (Fig. 5A), similar practice as described above was performed on 24-well plates. After 24 hours of incubation, the Matrigel chamber inserts were washed with phosphate buffered saline (PBS) on both sides, put back into wells that were filled with 0.75ml of Trypsin/EDTA Solution (0.05%), followed by the addition of another 0.5ml Trypsin/EDTA inside each insert. The trypsinized uninvaded cells (from above the Matrigel) were removed from interior of each Matrigel chamber insert into a new 15ml tube, and the trypsinized invaded cells (from below the membrane) were removed from each well into a new 15ml tube. Both cells were then washed with complete growth media and PBS before RNA was extracted.

### In vivo Metastasis Assays and Bioluminescence Imaging

Animal experiments described in this study were approved by West Virginia University (WVU) Institutional Animal Care and Use Committee and were performed at the WVU Animal Facility. 6-8 week-old female C57BL/6Ncrl mice were purchased from Charles River Laboratories and 6-8 week-old male NOD-scid IL2Rgamma^null^ (NSG, Stock No: 005557) mice were from The Jackson Laboratory. Generally, metastasis assays were performed with five mice in each group and replicated at least twice with independent cohorts, whereby similar results as described were achieved each time.

For experimental metastasis assays, mice were injected intravenously with indicated cells with *Luc2* expression. For B16F10 cells, 6×10^4^/mouse was used for NSG and 2×10^5^/mouse for C57BL/6Ncrl. Mice were euthanized on Day 15 post-injection for NSG and Day 21 post-injection for C57BL/6Ncrl. For YUMM1.7 cells, 1.5×10^5^/mouse was used for both NSG and C57BL/6Ncrl. Mice were euthanized on Day 24 post-injection for NSG and Day 32 post-injection for C57BL/6Ncrl. Lungs, livers and other organs (kidneys, brains, etc) were dissected and images were taken using an Olympus MVX10 Microscope. All organs were collected and frozen at -80°C for real time qPCR analysis. For tumor growth and spontaneous metastasis of B16F10 cells, C57BL/6Ncrl or NSG mice were injected subcutaneously with indicated cells with Luc2 expression (1.2X10^5^/mouse). Tumor volumes were recorded every other day from Day 7 or Day 8 post-injection to Day 21. All mice were then euthanized and organs were dissected for imaging and qPCR analysis.

Bioluminescence imaging was performed to quantify tumor burden *in vivo* one day before experimental animals were euthanized. Briefly, mice were injected intra-peritoneally with D-luciferin (Caliper Life Sciences, 150 mg/kg) and all images were taken between 10-20 minutes post-injection using the IVIS Lumina-II Imaging System (PerkinElmer, Waltham, MA) with 1.0 minute capture and medium binning. Living Image-4.0 software was used to process the captured images. Signal intensity was quantified as the sum of all detected photon counts within the region of interest after subtraction of background luminescence.

### Genomic DNA Extraction and Determination of Metastatic Tumor Burden

The method was described previously (42). It utilized two pairs of primers targeting firefly luciferase *Luc2* gene (only from injected tumor cells) and mouse *Ptger2* gene (from injected tumor cells and also mouse tissues) to calculate the relative ratio of metastatic melanoma cells within 10^4^ tissue cells. The number was used as a quantitative measurement of tumor burden in this work. Briefly, genomic DNA from mouse organs was extracted using Proteinase K digestion followed by ethanol precipitation. Each biological sample was then amplified in technical triplicate on a StepOnePlus Real-Time PCR System (Thermo Fisher Scientific) for Luc2 and Ptger2 fragments. On each plate, serial dilutions of B16F0-Luc2 or YUMM1.7-Luc2 genomic DNA were used for Luc2 and Ptger2 fragment amplification to create standard curves for the calculation of relative Luc2 DNA and total mouse DNA. Microsoft Excel 2013 was used to establish gene amplification standard curves (Ct vs. log DNA) for Luc2 and Ptger2. The relative Luc2 DNA amount (QLuc2) and total mouse (Ptger2) DNA amount (Qmm) for each genomic DNA sample were then calculated. The Luc2 cell ratio is calculated as: R = QLuc2 /Qmm. R is presented as Luc2 cell number in 10^4^ tissue (lung, liver, etc) cells.

### RNA Isolation and Gene Expression Analysis by Real Time qRT-PCR

All samples for RNA analysis were prepared in biological triplicates. Unless otherwise specified, all cells were plated on 6-well plates in complete growth medium for 48 hours before harvested for gene expression analysis. They may also be cultured in serum-free medium (SFM) for indicated time as described in the text before RNA extraction. Total RNA was isolated using GeneJET RNA Purification Kit (Thermo Fisher Scientific Inc., Waltham, MA) except for the invaded and uninvaded B16F10 cells in transwell assay (Fig. 5A), from which RNA was extracted using PicoPure RNA Isolation Kit (Thermo Fisher Scientific Inc). 50-500ng of RNA each was reverse transcribed using High Capacity RNA-to-cDNA Kit (Thermo Fisher Scientific). Real-time quantitative RT-PCR was performed on StepOnePlus Real-Time PCR System with Brilliant II SyBr Green QPCR Master Mix (Agilent Technologies, Santa Clara, CA). GAPDH served as the internal control for the reactions, and the normalized results were analyzed by GraphPad Prism (version 5). The primer pairs for indicated genes in the text were adopted from PrimerBank (58) and verified before assay use. The PrimerBank IDs will be provided upon request.

### Western Blotting Analysis

Whole-cell lysates for immunoblotting were obtained by extraction in ice-cold RIPA buffer (no SDS) with protease and phosphatase inhibitor cocktails (Thermo Fisher Scientific). Proteins were normalized by BCA assay (Pierce BCA Protein Assay Kit, Thermo Fisher Scientific), separated by SDS-PAGE, transferred to Bio Trace PVDF membrane (PALL Life Sciences, Pensacola, FL), probed by the indicated antibodies, and detected using SuperSignal West Pico PLUS Chemiluminescent Substrate (Thermo Fisher Scientific). Rabbit anti-WISP-1 (H-55, sc-25441) was from Santa Cruz Biotechnology (Dallas, Texas). The other rabbit polyclonal antibodies were purchased from Cell Signaling Technology (Danvers, MA): anti-β-actin (13E5), anti-Snail (C15D3), anti-ZEB1 (D80D3), anti-N-Cadherin (D4R1H), anti-Phospho-Akt (Ser473) (D9E), anti-Akt (pan) (C67E7), anti-Phospho-p44/42 MAPK (Erk1/2) (Thr202/Tyr204) (D13.14.4E) and anti-p44/42 MAPK (Erk1/2) (137F5).

### Cell Treatment with Conditioned Media, Recombinant Wisp1 and Kinase Inhibitors

To treat cells with conditioned medium containing overexpressed Wisp1 for EMT gene stimulation (Fig. 5G), B16F10-KO1 cells were seated on 6-well plates for 24 hours and grown in conditioned media from *Wisp1* -knockout NIH3T3-KO for another 24 hours. Thirty minutes before stimulation treatment, three groups of media were prepared. First group was conditioned media from *Wisp1* -knockout NIH3T3-KO, with antibody isotype control (Normal rat IgG, from Sigma-Aldrich, St. Louis, MO) at a final concentration of 20μg/ml. The second group was conditioned media from Wisp1 -overexpressed NIH3T3 -mWisp1, with the same antibody isotype control. The third group was conditioned media from Wisp1-overexpressed NIH3T3-mWisp1, with rat anti-Wisp1 (MAB1680, R&D Systems) at a final concentration of 20μg/ml. All three groups of media were incubated at 37°C for 30 minutes and then used to replace the media for B16F10-KO1 cells. The stimulation treatments were performed in biological triplicates for three hours and cells were harvested for RNA extraction and Real Time qRT-PCR analysis.

Recombinant mouse Wisp1 (rmWisp1, 1680-WS-050) was from R&D Systems and used at a final concentration of 5μg/ml following manufacturer’s instructions. Akt inhibitor MK-2206 (final 2.0 μg/ml) was from Sigma-Aldrich (St. Louis, MO) and MEK inhibitor U0126 (final 10 μM) was from Cell Signaling Technology (Danvers, MA). DMSO with the same volume was used for control cells. Unless otherwise specified, cell treatment for kinase immunoblot analysis maintained for 30 minutes while cell treatment for comparison of EMT marker gene expression maintained for 3 hours.

### Data Sets and Statistical Analysis

To compare melanoma with benign skin samples, the gene expression profiles of Affymetrix arrays were downloaded from Gene Expression Omnibus, GSE3189 (37). The abundance of WISP1 in primary melanoma and normal skin was quantified by immunohistological analysis using a tissue microarray derived from de-identified human skin tissue samples, as provided by the Human Protein Atlas (www.proteinatlas.org, Stockholm, Sweden (59)) and in accordance with approval from the Uppsala University Hospital Ethics Committee. The tissue microarray analysis included samples from 7 primary melanomas and 3 normal epithelial tissues that represented both male and female patients ranging in age from 46 to 87 years. The tissue microarrays were processed and analyzed as similarly described previously (35). In brief, processed tissue microarrays were probed using a rabbit polyclonal antibody against WISP1 (Sigma-Aldrich Cat #HPA007121, RRID:AB_1858844) that was validated by providing partly consistent staining patterns with previously reported gene/protein data, weak band of predicted size in western blot validation, and passing protein array validation tests. WISP1 staining was visualized using diaminobenzidine and microscopic tissue features were visualized by counterstaining with Harris hematoxylin. Immunohistochemically stained tissue microarrays were scanned at 20x resolution (1 mm diameter) and provided as an 8-bit RGB JPEG image. The average intensity of WISP1 staining per tissue sample was quantified by deconvoluting the intensity of WISP1 staining from nonspecific hematoxylin tissue staining in R using the EBImage package. Following color deconvolution, the image was segmented into tissue and non-tissue regions. A tissue mask used for segmenting IHC images was determined based on non-zero staining in any of the RGB channels, following background image correction. To address the question of whether more of the TMA image stains positive for WISP1 (i.e., there are more cells that produce WISP1 within the tissue sample) but that the intensity of WISP1 staining is the same (i.e., WISP1 production per cell is not increased) in melanoma samples, we calculated the distribution in WISP 1 staining intensity and the fraction of the total tissue area that stains strongly for WISP1. To compare the gene expression profiles of WISP1 in primary melanoma with overall survival, Level 3 skin cutaneous melanoma (SKCM) RNAseqV2 mRNA expression results [FPKM normalized] and clinical profiles for patients diagnosed with primary melanoma that had not metastasized (i.e., stage I to III) were obtained from the Cancer Genome Atlas (TCGA). Individual statistical methods are indicated in the figure legend.

Unless specified, all analyses were performed with GraphPad Prism (version 5). Individual quantitative result was shown as Mean±SEM. Box plots indicate median and inter-quartile range (box), 5th and 95th percentiles (whiskers). Data sets were compared using the unpaired Student’s t-test (two-tailed) or one-way analysis of variance (ANOVA) followed by Tukey’s multiple comparison *ad hoc* post-test. To estimate cumulative survival probability, Kaplan-Meier survival curves were estimated from the cohort overall survival data. Statistical significance associated with a difference in survival between two groups was estimated using the Peto & Peto modification of the Gehan-Wilcoxon test and the Cox proportional hazards regression model, as implemented in the R REFERENCES *survival* package. A p-value of <0.05 was considered statistically significant. Asterisks are used to indicate the numerical value, where *: p-value < 0.05; **: p-value < 0.01; ***: p-value < 0.001; and ns: indicates not significant.

## ACKNOWLEDGMENTS

This work was supported by National Science Foundation (NSF CBET-1644932 to DJK) and National Cancer Institute (NCI 1R01CA193473 to DJK). The content is solely the responsibility of the authors and does not necessarily represent the official views of the NSF or NCI. Small animal imaging and image analysis were performed in the West Virginia University Animal Models & Imaging Facility, which has been supported by the West Virginia University Cancer Institute and NIH grants P20 RR016440 and P30RR032138/P30GM103488.

## CONFLICT OF INTEREST

The authors declare that they have no conflicts of interest with the contents of this article.

## AUTHOR CONTRIBUTIONS

Conceptualized study: WD and DJK; performed experiments: WD, AF, and SLM; analyzed data: WD, AF, and DJK; and drafted original manuscript: WD and DJK. All authors edited and approved of the final version of the manuscript.

---

**Supplementary Table S1.** Summary of mutation rate that includes both changes in single nucleotides and in copy numbers for TCGA melanoma (SKCM) samples from where both types of mutational data are reported. Gene mutation statistics from indicated numbers of cutaneous melanoma samples (SKCM) was obtained from the Cancer Genome Atlas (TCGA) through cBioPortal (retrieved 09/24/2018 - 52,53). Q-value corresponds to the probability of the observed or more extreme copy number variant is explained by random chance, where the value has been adjusted using the Benjamini and Hochberg FDR correction.

| Gene | Melanoma samples with Single Nucleotide Variant (SNV) and Copy Number Variant (CNV) data (N=287) |  | Q-value for CNV |
| --- | --- | --- | --- |
|  | Rate | Type |  |
| **NRAS** | 30% | 87 SNV 8 Amplification | 9.79E-07 |
| **BRAF** | 51% | 143 SNV 18 Amplification | 1.64E-07 |
| **PTEN** | 15% | 25 SNV 19 Deletion | 1.85E-27 |
| **TP53** | 17% | 48 SNV 3 Deletion | 3.69E-02 |
| **CDKN2A** | 43% | 40 SNV 1 Amp 88 Del | <1.0e-30 |
| **CTNNB1** | 7% | 16 SNV 3 Amplification | 1.39E-01 |
| **MITF** | 8% | 6 SNV 19 Amplification | 2.27E-13 |
| **KIT** | 7% | 12 SNV 10 Amplification | 2.39E-03 |
| **WISP1** | 8% | 7 SNV 16 Amplification | 2.90E-05 |

**Supplementary Table S2.** Wisp1 secretion from different cell lines cultured in 2D. Relative Wisp1 concentration (CONC), presented as mean with standard error, was determined for 48-hour conditioned medium of each cell line plated at a similar confluence with ELISA using human recombinant WISP1 as standard. n.d., not detected.

| Cell | CONC (pg/ml) | Cell | CONC (pg/ml) |
| --- | --- | --- | --- |
| B16F0 | 605 ± 15 | B16F10-KO1 | n.d. |
| B16F10 | 1,300 ± 35 | B16F10-KO2 | n.d. |
| E0771 | < 20 | B16F10-KO1-pBabe | n.d. |
| LLC1 | < 20 | B16F10-KO1-hSnai1 | n.d. |
| Melan-A | 1018 ± 32 | B16F10-KO1-mWisp1 | 3465 ± 233 |
| NIH3T3 | 83 ± 2.2 | NIH3T3-KO | n.d. |
| YUMM1.1 | < 20 | NIH3T3-pBabe | 90.3 ± 2.8 |
| YUMM1.7 | 451 ± 25 | NIH3T3-mWisp1 | 920 ± 15.6 |
| RPMI-7951 | 1,331 ± 34 | YUMM1.7-KO1 | n.d. |
| SK-MEL-3 | n.d. | YUMM1.7-KO2 | n.d. |
| SK-MEL-24 | n.d. | RPMI-7951-KO1 | n.d. |
| SH-4 | n.d. | RPMI-7951-KO2 | n.d. |

**Supplementary Figure S1.**
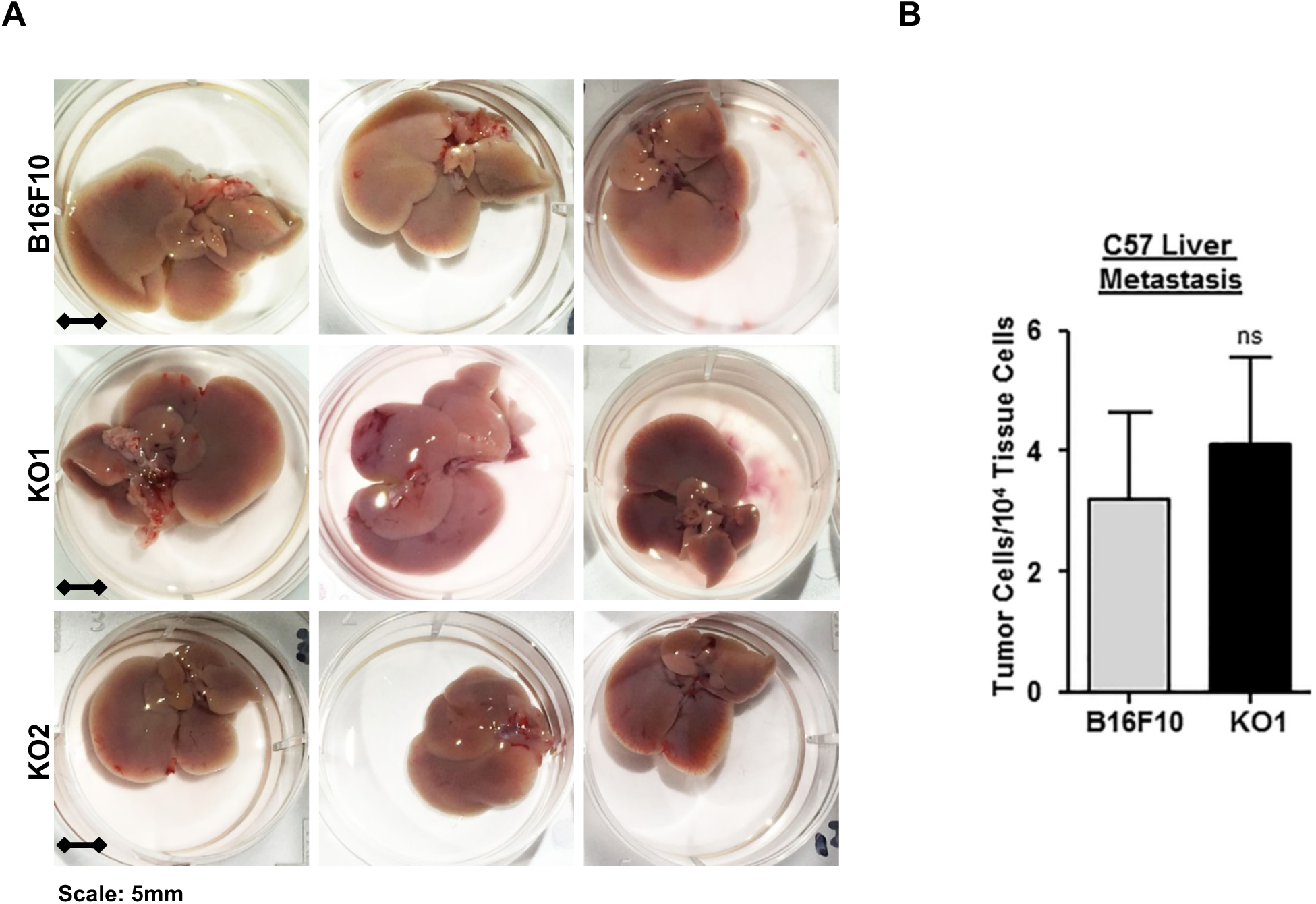
No difference was observed on livers of C57BL/6Ncrl mice from experimental metastasis assay using B16F10 and its knockout cells. (A) Representative livers from C57BL/6Ncrl mice with B16F10 or knockout cells (-KO1 and -KO2) in experimental metastasis assay (**Fig. 3G-3I**). (B) Real time genomic qPCR quantitatively comparing tumor liver metastatic burdens (tumor cell number within 10,000 mouse tissue cells). ns, not significant.

**Supplementary Figure S2.**
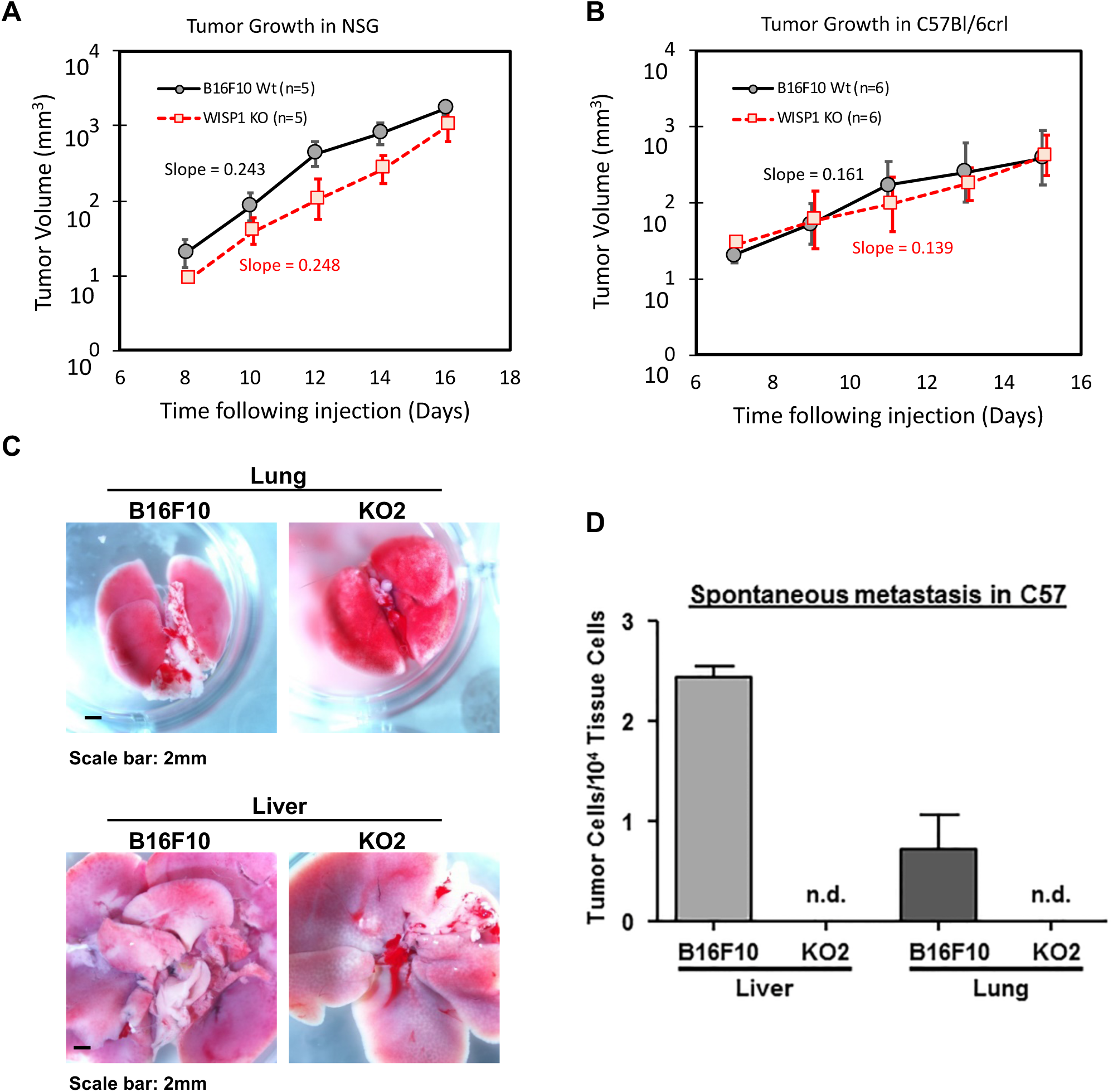
Real time genomic qPCR revealed that Wisp1 knockout repressed the spontaneous metastasis of melanoma cell line B16F10 in C57BL/6Ncrl mice. Growth of tumors derived from B16F10 or its knockout cell (-KO2) were monitored following subcutaneous injection in NSG mice (A: B16F10 (n = 5) or WISP1 KO cell (n = 5)) or in C57BL/6Ncrl mice (B: B16F10 (n=6) or WISP1 KO cell (n=6)). After 21 days, remaining C57BL/6crl mice (n = 4 in each group) were euthanized and lungs and livers were assayed for B16F10 tumor cells using real time genomic qPCR, as described in Materials and Methods. (C) Representative lungs and livers from C57BL/6Ncrl mice with B16F10 or knockout cell at day 21. (D) Real time genomic qPCR results showed quantitative tumor lung and liver metastatic burden in spontaneous metastasis assays, n.d., not detected.

